# Interspecies hybridization as a route of accessory chromosome origin in fungal species infecting wild grasses

**DOI:** 10.1101/2024.10.03.616481

**Authors:** Wagner C. Fagundes, Mareike Möller, Alice Feurtey, Rune Hansen, Janine Haueisen, Fatemeh Salimi, Alireza Alizadeh, Eva H. Stukenbrock

## Abstract

Many fungal plant pathogens have dynamic genomic architectures that can contribute to rapid evolution and adaptation to new niches. *Zymoseptoria tritici*, an important fungal pathogen of wheat, has a compartmentalized and rapidly evolving genome. In the genome of the reference isolate *Z. tritici* IPO323, 8 of the 21 chromosomes are accessory. In spite of the profound impact on genome organization, the origin of accessory chromosomes in *Z. tritici* is still poorly understood. Combining genomics, transcriptomics and epigenomics, we discovered a new chromosome in *Z. tritici* isolates infecting wild grasses from the genus *Aegilops*, and we use this discovery to study the origin of accessory chromosomes. The newly identified chromosome presents similar characteristics to known accessory chromosomes in *Zymoseptoria* species, including presence-absence variation, low gene expression *in vitro* and *in planta*, and enrichment with heterochromatin-associated histone methylation marks (H3K27me3). Interestingly, we found an orthologous chromosome in *Zymoseptoria ardabiliae*, a closely related fungal species also infecting wild grasses. This ortholog chromosome also presents accessory chromosomes characteristics, but lacks the enrichment of heterochromatin-associated methylation marks. Transcriptomic analyses revealed that the orthologous chromosome in *Z. ardabiliae* harbors active transposable elements (TEs) congruent with lower signatures of host-genome defense mechanisms against TE expansion and spread (quantified as repeat-induced-point (RIP) mutation signatures). Our findings suggest that the chromosome has been exchanged between *Z. tritici* and *Z. ardabiliae* by introgressive hybridization events underlining the relevance of hybridization in the evolution of new accessory chromosomes. We speculate that the regulation of TEs has not yet occurred on this new accessory chromosome in *Z. ardabiliae*, contributing to its rapid evolution.

## Introduction

Many fungal plant pathogens exhibit remarkable variation in genome architecture, even between closely related species (1). Key observations from comparative genome studies involved the identification of distinct and rapidly evolving compartments in genomes of closely related fungal plant pathogen species, including gene-sparse, repeat-rich regions, AT-rich isochores, clusters of tandem duplicated genes and accessory or lineage-specific chromosomes (2–11). These distinct genome compartments present different rates of mutation and recombination events and contribute to dynamic genome architectures and rapid evolution of fungal pathogens (1, 7, 12).

Fungal accessory chromosomes are generally small (< 2Mb in length), enriched with Transposable Elements (TEs) and have a lower gene density and transcriptional activity compared to core chromosomes (13, 14). Moreover, accessory chromosomes were experimentally shown to be lost recurrently during mitotic cell divisions and to undergo non-mendelian segregation in the fungus *Zymoseptoria tritici*, resulting in a presence-absence variation (PAV) among individuals of a population (15, 16). In *Z. tritici* as well as in other species, accessory chromosomes were also shown to be enriched in heterochromatin-associated histone methylation marks (e.g. H3K27me3 and H3K9me3), correlating with the low transcriptional activity found on these chromosomes (13, 14, 17). In some fungal species, accessory chromosomes encode virulence-related genes that can determine disease outcome and host range of specific fungal species or lineages, as reported for *Leptosphaeria maculans* (18), *Fusarium solani* (19) and *Fusarium oxysporum* f. sp. *lycopersici* (3, 20). Despite the PAV between fungal individuals, accessory chromosomes appear to have been maintained over long evolutionary periods, raising questions on the transmission and maintenance of chromosomes over generations and more importantly, how they originate in the first place. Few studies have experimentally addressed these questions in different fungal pathogens (3, 21, 22), but the evolutionary mechanisms of accessory chromosome-origin remain largely unknown.

*Zymoseptoria tritici*, besides being an important fungal pathogen of wheat, is also an important experimental model in studies of fungal genome evolution (23, 24). Coalescence analyses suggest that emergence of this wheat pathogen from wild ancestors occurred in the Fertile Crescent region in the Middle East and coincided with the domestication of wheat (25). Population genetics studies using polymorphism data and whole genome sequencing from worldwide *Z. tritici* populations have indicated high levels of genetic variation within and between *Z. tritici* populations and extensive gene flow from regional to continental scales (26–29). The reference genome of *Z. tritici* is one of best assembled haploid fungal genomes with 21 chromosomes sequenced from telomere to telomere in the isolate IPO323, eight of them being accessory (28). Accessory chromosomes in *Z. tritici* show hallmarks that distinguish them from the core chromosomes: whole chromosome PAV between isolates; TE-richness; low gene density; enrichment of the heterochromatin-associated methylation mark H3K27me3; and low gene transcriptional activity not only *in vitro* but also *in planta* (17, 21, 28, 30–35). In contrast to other fungal pathogens, accessory chromosomes in *Z. tritici* are not enriched in virulence-related genes and no virulence factors have yet been identified on these chromosomes, even though findings suggest that they can confer a fitness cost in a wheat cultivar-specific manner (36).

The closest known relatives of *Z. tritici* namely *Z. brevis*, *Z. pseudotritici* and *Z. ardabiliae* are found to be associated with different wild grasses in the Middle East as e.g. *Lolium* spp., *Elymus repens*, and *Dactylis glomerata* (37–39). Comparative genomics analyses using long-read assemblies and gene annotations of different *Zymoseptoria* species have revealed a diverse set of genes and a large distribution of accessory chromosomes within and between species, suggesting that genome compartmentalization is an ancestral trait in the *Zymoseptoria* genus (29, 40). Comparative analyses of genome composition across different *Zymoseptoria* species revealed a prominent role of interspecific hybridization in species evolution. Notably, at the center of origin of these pathogens, where they co-exist, hybridization appears to occur readily and to shape patterns of genetic variation (41–45). Intriguingly, hybridization can also confer adaptive introgression of functionally relevant traits as demonstrated for the DNA methyltransferase gene *DIM2* (44). However, to which extent hybridization impacts accessory chromosome evolution has so far not been addressed.

Recently, we have identified host-specific populations of *Z. tritici* infecting wild grass species of the genus *Aegilops*. Preliminary analyses indicate that *Aegilops*-infecting *Z. tritici* isolates are distinct from wheat-infecting *Z. tritici* isolates and from closely related *Zymoseptoria* species (46). Different footprints of positive selection were detected between host-diverging populations and demography analyses further suggested a split of the wheat- and *Aegilops*-infecting lineages after 10,000 years ago (46). The collection of closely related lineages of *Z. tritici* provides an excellent model system to address the evolution of genome architecture and accessory chromosome composition in natural fungal populations.

In this study, we explore the genomic variation in *Z. tritici* isolates infecting *Aegilops* species. We present a new high-quality reference genome based on long read-sequencing for the *Aegilops*-infecting *Z. tritici* Zt469 coupled with gene and Transposable Elements (TEs) predictions, and compare the gene content between this isolate and other *Z. tritici* isolates as well as between closely related *Zymoseptoria* species. Combining different omics’ approaches, we identify a unique accessory chromosome in *Aegilops*-infecting *Z. tritici* isolates which has synteny to another accessory chromosome in the closely related *Z. ardabiliae* species. Analyses of the orthologous chromosomes in the two *Zymoseptoria* species reveal different levels of TE expression and host-genome defense mechanism activity against TEs besides distinct enrichment of heterochromatin-associated methylation marks. We suggest that this chromosome has been exchanged between *Z. tritici* and *Z. ardabiliae* and that interspecies hybridization combined with TE activity can play a prominent role in the evolution and diversification of new accessory chromosomes.

## Material and methods

### DNA sequencing and whole-genome assemblies

Reference genomes for *Aegilops*-infecting *Z. tritici* isolate Zt469 and *Z. ardabiliae* isolate Za100 were sequenced using the single-molecule real-time (SMRT) PacBio technology. Zt469 was isolated from *Aegilops* spp. in 2018 and the *Z. ardabiliae* isolate Za100 was isolated in 2011 from *Agropyrum tauri*. High quality DNA was extracted as described previously (47). Library preparations and sequencing were performed at the Max Planck-Genome-Centre, Cologne, Germany using a PacBio Sequel II platform. Genomes were assembled *de novo* using the SMRT Analysis software v.5 (Pacific Bioscience) using default and “fungal” parameters as previously described (29). The genome assemblies with the best quality determined by the software Quast (48) were used for further analysis. A previous version of the Zt469 genome assembly has been published (44) and an improved version of this genome assembly is presented here. In this version, we have filtered the Zt469 assembly to match filtering steps done previously and to improve the overall genome assembly quality (10, 29); Supplementary Table S1). Telomeric repeats for each contig were identified as previously described (29); Supplementary Table S1). Genome assemblies and gene annotations based on long-read sequencing (PacBio) of the *Z. tritici* isolates Zt05 and Zt10 and of the closely related species *Zymoseptoria brevis* (Zb87), *Z. passerinii* isolates (Zpa63 and Zpa796), *Z. pseudotritici* (Zp13) and *Z. ardabiliae* (Za17) were obtained from previous studies (29, 49). Genome assembly, gene and repeat annotations from the reference wheat-infecting *Z. tritici* isolate IPO323 were also obtained from previous studies (28, 40, 45). Population data from *Aegilops*- and wheat-infecting *Z. tritici* and from additional *Z. ardabiliae* isolates were also obtained from elsewhere (38, 41, 46, 50).

### Repeat and gene annotations

Gene and TE annotations for Zt469 and Za100 genomes was done using a previously published pipeline (29). In brief, TE content in both genomes was annotated using the TEdenovo and TEannot tools from the REPET package (https://urgi.versailles.inra.fr/Tools/REPET; (51, 52) following the developer’s recommendations and default parameters as described in Feurtey et al. (29). TEs were classified according to the nomenclature defined by Wicker et al. (53). For visualization in Circos plots (54), we calculated gene and TE densities in 100 kb non-overlapping windows using bedtools v2.26.0 (55).

We used distinct tools to predict the putative gene functions in the newly assembled Zt469 and Za100 genomes. First, we obtained the amino acid sequence of each gene model and used the tool Predector to predict secreted proteins and PFAM domains (56). Predector uses several tools for fungal secretome and effector analyses, including the softwares SignalP (versions 3, 4, 5, 6; (57–60), TargetP (version 2.0; (61), DeepLoc (62), TMHMM (63, 64), Phobius (65), DeepSig (66), CAZyme finding (with dbCAN; (67), Pfamscan (68), searches against PHI-base (69), Pepstats (68), ApoplastP (70), LOCALIZER (71), Deepredeff (72), and EffectorP (versions 1, 2 and 3; (73–75). We furthermore used the software eggnog-mapper to provide additional Clusters of Orthologous Groups (COG), GO (Gene Ontology) and KEGG (Kyoto encyclopedia of genes and genomes) annotations (76). Then, in order to analyze gene expression, we categorized the gene models into “Effector” and “CAZyme” based on the individual tools score thresholds recommended by Predector (Supplementary Table S2). “Small Secreted Proteins (SSPs)” category also followed the criteria described previously (40); Supplementary Table S2). At last, we used Antismash v.6.0 (fungal version) to detect biosynthetic gene clusters (BGCs) in each genome (77); Supplementary Table S2). Gene models that did not belong to any of these categories were classified as “Other”. The outputs of all tools and transcription *in vitro* and *in planta* (when applicable) for each gene model as well as the genes belonging to each category can be found at Supplementary Tables S3-S5.

### Synteny and orthologous genes analyses

We identified orthologous genes between the *Zymoseptoria* genome assemblies using the software PoFF (78) implemented in Proteinortho (79) to account for synteny information. The identified orthogroups were used to visualize synteny between genomes using the software Circos (54).

### ChIP- and RNA-seq data analyses

Preparation and sequencing of RNA- and ChIP-seq samples of *in vitro* growth are described in the Supplementary Text S1. For RNA-seq *in planta*, we analyzed infection stage-specific transcriptome data generated in a previous study (46). Sequencing adapters and low-quality reads were trimmed from raw paired-end RNA and ChIP-seq reads using Trimmomatic v 0.39 (80) with the following parameters: LEADING:20 TRAILING:20 SLIDINGWINDOW:5:20 MINLEN:50. Low quality nucleotides (Q<20) in RNA-seq reads were further masked using FASTX-toolkit v0.0.13 (http://hannonlab.cshl.edu/fastx_toolkit/). For RNA-seq data, quality-trimmed and masked reads were mapped to the Zt469 and Za100 genomes using HISAT2 (81) while for ChIP-seq read mapping was performed using Bowtie2 (82). Conversion, sorting, merging and indexing of alignment files were performed using SAMtools v. 1.7 (83). We used the software HOMER (84) to detect methylation-enriched regions in the ChIP mappings. We called methylation-enriched peaks individually for each replicate and merged the peaks with bedtools v2.26.0 (55). Only enriched regions found in all replicates were considered for downstream analyses. Sequence coverage of methylation-enriched regions per contigs were also calculated using bedtools v2.26.0 (53). For RNA-seq *in vitro* and *in planta*, we accessed gene expression as Transcript Per Million (TPM) as previously described (29). For Circos plot representations (54), we first calculated the mean TPM between replicates and then the averages per 100 kb non-overlapping windows in each chromosome/unitig based on the log2(TPM+1) values. For RNA-seq read mapping statistics, we used the tool Qualimap bamQC v.2.2.1 (85).

### Transposable Element expression analysis

We evaluated Transposable Elements (TEs) transcription activity in both Zt469 and Za100 genomes using the *in vitro* and *in planta* RNA-seq data. Raw RNA-seq reads were trimmed, quality filtered and masked as described above. Quality-trimmed and masked reads were mapped to each respective genome assembly using HISAT2 (81) with the flag “--no-mixed”. Conversion, sorting, merging and indexing of alignment files was also performed using SAMtools v. 1.7 (83). Then, we accessed the TE transcription *in vitro* and *in planta* for each replicate and each infection stage individually. To this end, read count tables were generated using the TEcount function of TEtranscript pipeline with the “--mode multi” (86, 87) and levels of expression were reported using the Transcript per Million (TPM) normalization as described previously (29, 44). The length of each TE was calculated with the GenomicFeatures R package using the function “exonsBy”, in which the entire element was treated as a unique exon (88).

### Repeat-induced point mutation (RIP) analyses

We quantified RIP signatures along the Za100 and Zt469 genomes and among TE copies within each genome. To this end, RIP-like signatures and the Large RIP-affected genomic regions (LRARs) were calculated using the RIPper software (89). Genome-wide RIP composite indices were calculated using default parameters of 1000bp windows with a 500bp step size and following the formula: (TpA/ApT) – (CpA þ TpG/ApC þ GpT) (88). LRARs were defined as genomic regions (windows) consecutively affected by RIP that are more than 4000bp in length. To access the RIP composite index of each TE copy, we calculated the indices in 50bp nonoverlapping windows using a previously published custom script (45). Regions are considered to be affected by RIP when the composite index is > 0 (88).

### Karyotype analysis

We compared the genome structure and chromosome presence-absence variation (PAV) of different *Z. tritici* and *Z. ardabiliae* isolates using two approaches. Firstly, whole-genome sequencing reads were trimmed using Trimmomatic v 0.39 (80) using the following parameters: LEADING:20 TRAILING:20 SLIDINGWINDOW:5:20 MINLEN:50. Trimmed reads were then mapped to the *Z. tritici* reference genome IPO323 (28) as well as to the newly assembled *Aegilops*-infecting *Z. tritici* Zt469 and *Z. ardabiliae* Za100 genomes using bwa-mem v. 0.7.17 (90). After mapping, we normalized the coverage of mapped reads by each assembly size using deepTools v.2.0 (91). Normalized alignment files were then visualized with the Integrate Genome Viewer (IGV) software v.2.8.2 (92). Only the Zt469 and Za100 contigs containing telomeric repeats and/or larger than 100 kb were kept for visualization. A second approach involved the analysis of chromosome PAV by Pulsed Field Gel Electrophoresis (PFGE) and Southern blot techniques (93). Preparation of non-protoplast plugs for PFGE analyses was done as described previously (35). As a comparison to Zt469, we also prepared non-protoplast plugs for PFGE analyses for the reference *Z. tritici* IPO323 isolate (strain Zt244) (28), the wheat-infecting *Z. tritici* isolate Zt10 (strain Zt366; (25) and the *Aegilops*-infecting *Z. tritici* isolate Zt501. PFGE running conditions and Southern blot analyses were done as previously described and are further summarized in the Supplementary Text S1.

### Introgression and nucleotide diversity analyses

We calculated the Patterson’s *D*-score, also known as the ABBA-BABA test (94, 95), to determine the occurrence of interspecific gene flow between *Z. tritici* and *Z. ardabiliae*. The test is based on a resolved phylogeny among four taxa (P1, P2, P3, O) and determines, along the genome, the proportion of derived (‘B’) and ancestral (‘A’) alleles as defined by the outgroup (O). An excess of shared derived alleles indicates gene flow between two of the taxa (92, 93). In our tests, we placed either the *Aegilops*-infecting *Z. tritici* isolate Zt469 or the *Aegilops*-infecting *Z. tritici* isolate Zt436 in position P1; the wheat-infecting *Z. tritici* isolate Zt565 or the wheat-infecting isolate Zt668 in position P2; and either the *Z. ardabiliae* isolate Za100 or Za98 in position P3 (Supplementary Figure S1). We performed the tests with two different *Aegilops*-infecting *Z. tritici* isolates and two different *Z. ardabiliae* isolates to check if gene flow would occur only in isolates carrying the unitig 9 (in the case of Zt469) or unitig 3 (in the case Za100). For all tests, we used the *Z. passerini* Zpa796 isolate as the outgroup (O). A negative *D* statistic in these configurations is indicative of gene flow between *Aegilops*-infecting *Z. tritici* (P1) and *Z. ardabiliae* (P3) while positive scores indicate gene flow between wheat-infecting *Z. tritici* (P2) and *Z. ardabiliae* (P3). Details of genome data filtering and processing for ABBA-BABA tests are provided in the Supplementary Text S1.

For the nucleotide diversity analyses in the syntenic unitig 9 and unitig 3 chromosomes, we used Single Nucleotide Polymorphisms (SNPs) obtained from population data of *Aegilops*-infecting *Z. tritici* and *Z. ardabiliae* isolates. The complete description of whole-genome sequencing read trimming, mapping and variant calling is described on the Supplementary Text S1. From the resulting VCF files, we calculated nucleotide diversity (π) (96) in 1 kb windows using VCFtools v 0.1.13 (97) with the haploid mode fork provided by Julien Y. Dutheil (https://github.com/vcftools/vcftools/pull/69). We compared nucleotide diversity between unitig 9 in Zt469 and unitig 3 in Za100 using polymorphism data from three *Aegilops*-infecting *Z. tritici* and the three *Z. ardabiliae* isolates that contained these chromosomes as identified by the read-mapping approach. The three *Aegilops*-infecting *Z. tritici* isolates were selected based on the lowest pairwise Identity-By-State (IBS) similarity values acquired using PLINK v. 1.07 (98) and the generated VCF file in order to keep the most genotypically diverse isolates.

## Results

### A new accessory chromosome in *Aegilops*-infecting *Z. tritici*

We first used high-quality genome data from *Aegilops* and wheat-infecting *Z. tritici* isolates to characterize the extent of structural conservation between the two host-specific lineages. Analyses of homologous genes between the reference wheat-infecting *Z. tritici* isolate IPO323 (28) and the *Aegilops*-infecting *Z. tritici* isolate Zt469 revealed a high extent of synteny between the thirteen core chromosomes described for IPO323, whereas only three out of eight accessory chromosomes were found to have synteny with chromosomes of Zt469 (unitigs 18, 19 and 20) (Figure 1). Unexpectedly, this analysis also revealed a complete chromosome (with telomeric repeats on both ends) in Zt469 with no alignments to any specific chromosome of the reference strain IPO323, namely “unitig 9” (Figure 1). Only 4.3% (13/299) of the precited genes on unitig 9 have homologs in the IPO323 genome and the 13 genes are distributed across different chromosomes of IPO323.

**Figure 1.**
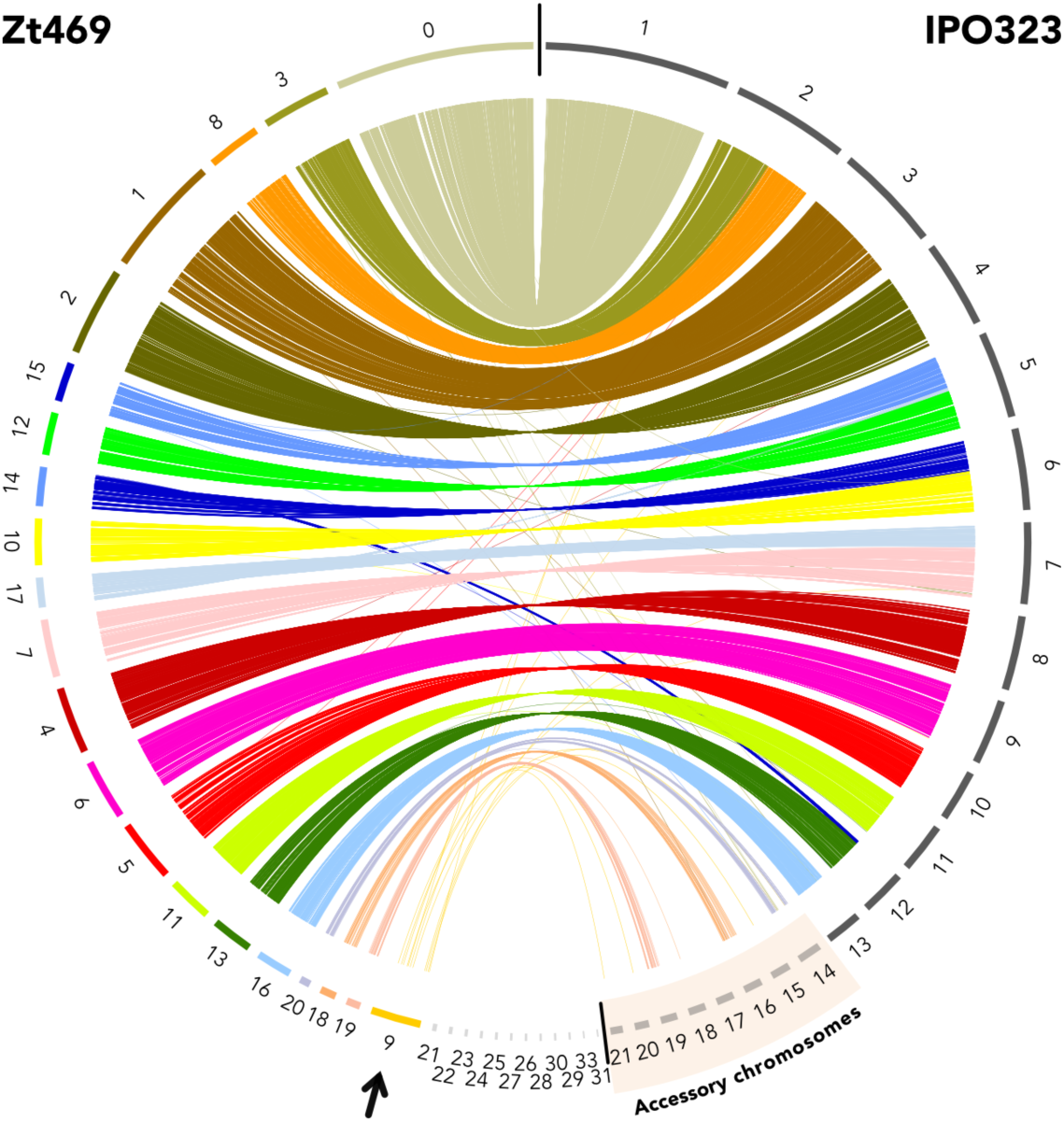
Genome synteny between *Z. tritici* isolates Zt469 and IPO323. Synteny analysis was performed between the reference wheat-infecting *Z. tritici* isolate IPO323 (right) and the *Aegilops*-infecting *Z. tritici* isolate Zt469 (left). Each color represents a different unitig in Zt469 and colored lines represent homologous genes. Small unitigs are shown in light gray. A new chromosome, referred to as “unitig 9”, is identified in Zt469 and show no synteny to any particular portion of the IPO323 genome (black arrow). Unitigs in Zt469 are ordered following the synteny to the reference IPO323 genome. Accessory chromosomes on the IPO323 genome are indicated.

To further validate the correctness of our genome assembly and the presence and size of unitig 9 in Zt469, we conducted Pulsed Field Gel Electrophoresis (PFGE) followed by Southern blot analysis, which confirmed the presence of a chromosome in the expected size (∼1.6Mb) in Zt469 and the absence of the chromosome in the *Z. tritici* IPO323 isolate and in the other two randomly selected *Z. tritici* isolates analyzed (Supplementary Figure S2).

### Unitig 9 exhibits presence-absence variation in *Aegilops*-infecting *Z. tritici* isolates

In order to examine the presence of unitig 9 in other sympatric *Z. tritici* isolates collected in the Middle East, we conducted genome-wide comparative analyses to identify chromosome presence-absence variation (PAV) between Iranian *Z. tritici* isolates infecting leaves of wheat and *Aegilops* spp. collected in agricultural fields and natural grasslands, respectively. First, using the IPO323 reference genome and a normalized read coverage approach, we find extensive PAV of the accessory chromosomes for all analyzed *Z. tritici* isolates (Supplementary Figure S3). Hereby, we also find that the wheat-infecting isolates exhibited an overall higher presence of these chromosomes when compared to the *Aegilops*-infecting ones (Supplementary Figure S3). Next, we used the same mapping approach based on the *Aegilops*-infecting Zt469 reference genome. Hereby, we observe a similar pattern of chromosome PAV for the three shared accessory unitigs (unitigs 18, 19 and 20) while unitig 9 was completely absent in all wheat-infecting isolates (Figure 2). When analyzing the *Aegilops*-infecting population, we observe however that unitig 9 was present in 11 out of 47 isolates while the accessory unitigs 18, 19 and 20 occurred more frequently in the population (among the 47 isolates in 38, 24 and 23 isolates, respectively) (Figure 2). These results indicate that unitig 9 is a chromosome present only in *Aegilops*-infecting *Z. tritici* populations from the Iranian pathogen population.

**Figure 2.**
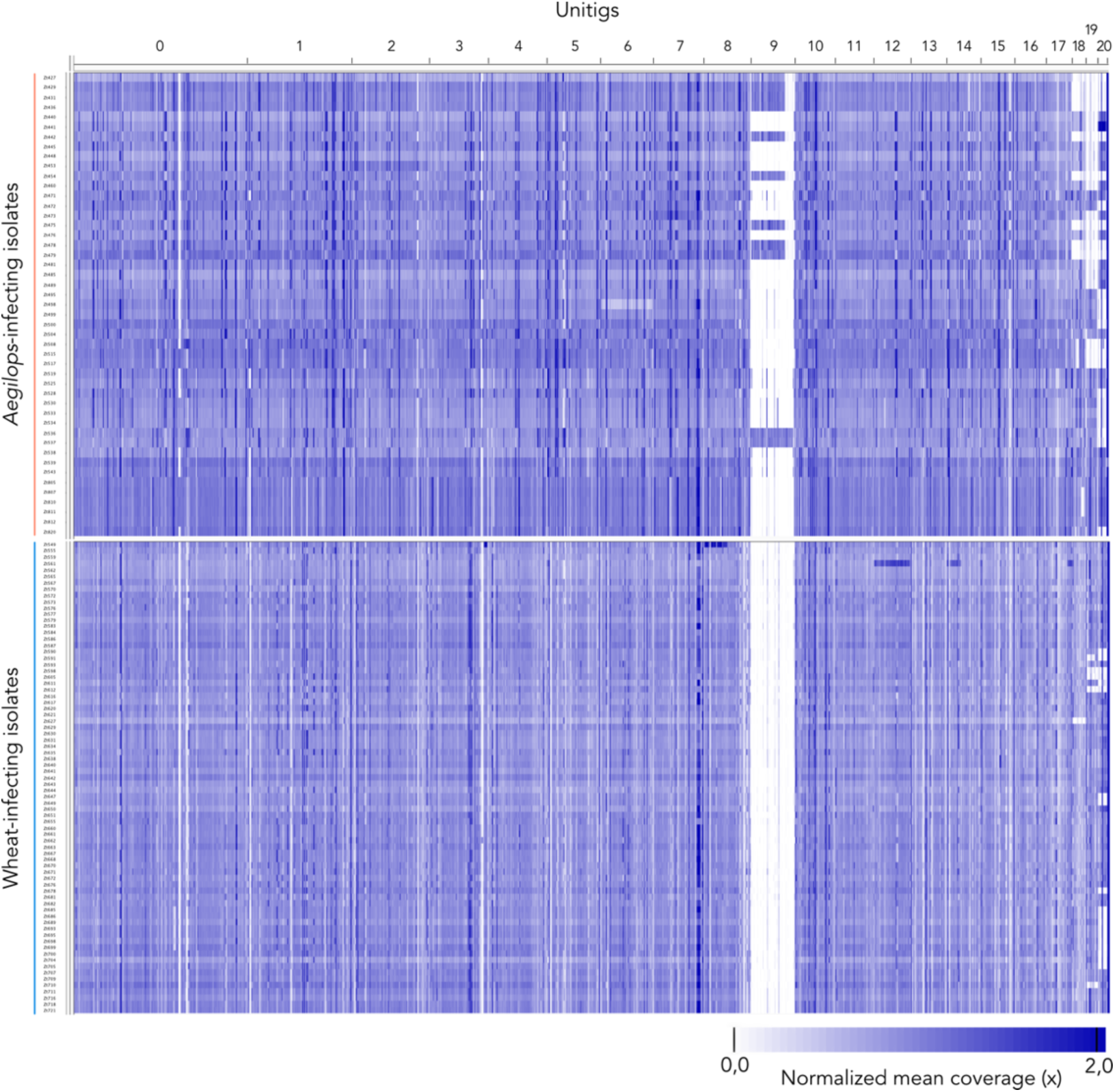
Unitig 9 shows presence-absence variation in *Aegilops*-infecting *Z. tritici*. Chromosome presence-absence variation (PAV) in host-diverging *Z. tritici* populations was analyzed by read mapping to the *Z. tritici* Zt469 genome assembly. The heatmap represents normalized mean coverage of reads mapped to each position of the genome assembly. Except for unitig 9, only unitigs syntenic to the *Z. tritici* IPO323 reference genome are shown and sorted in descending order of length. Darker colors represent regions of higher coverage as e.g. repetitive elements or duplications.

### Unitig 9 exhibits accessory chromosome hallmarks

Considering the PAV of unitig 9 among *Aegilops*-infecting *Z. tritici* isolates, we hypothesized that unitig 9 represents a new accessory chromosome. We therefore set out to investigate if unitig 9 shares the same hallmarks described for accessory chromosomes in *Zymoseptoria* species as e.g. low gene density; high TE density and low transcriptional activity (1, 28, 29). Gene and TE annotations as well as transcriptome analyses *in vitro* indeed revealed that unitig 9 has a lower gene density (average of 0.39 genes / kb), higher TE density (average of 0.33 TEs / kb) and lower transcriptional activity when compared to core unitigs of similar size (e.g. unitig 8; 0.83 genes/kb and 0.28 TEs/kb on average) (Figure 3A), consistent with known signatures of other accessory chromosomes in *Z. tritici* (28, 31). The smaller unitigs 18, 19 and 20 which we also consider as accessory chromosomes exhibited similar characteristics (Figure 3A).

**Figure 3.**
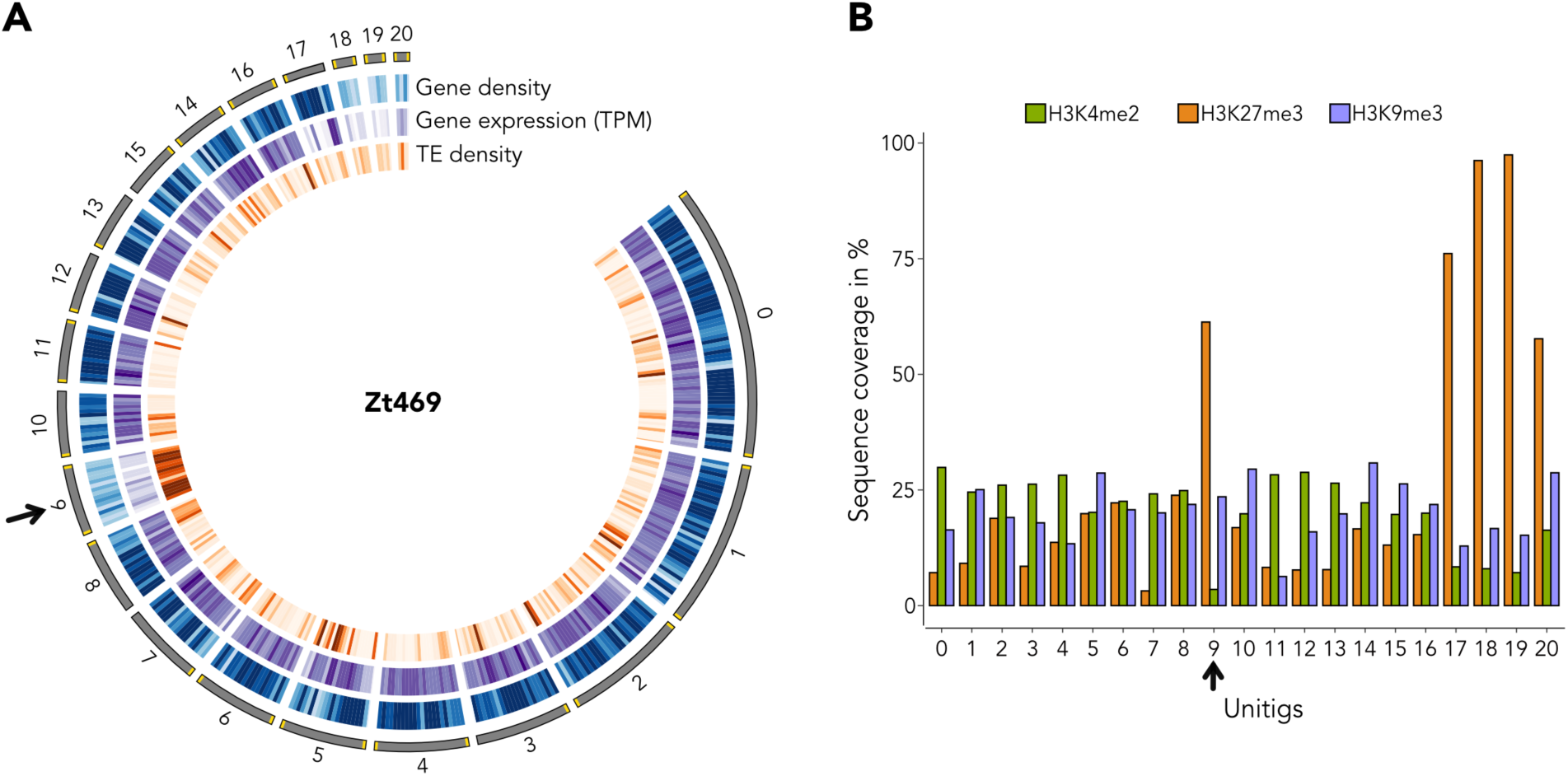
Unitig 9 presents accessory chromosome characteristics. **(A)** Circos plot representing different features along twenty Zt469 unitigs. Unitigs are represented by the dark gray segments sorted by length. Telomeric repeats are indicated in yellow. Tracks from outside to the inside are heatmaps in 100kb windows representing respectively: gene density, gene expression *in vitro* in log2(TPM+1) and TE density. Darker heatmap colors indicate higher values. Unitig 9 is indicated by a black arrow. **(B)** Barplot displaying the percentage of sequence coverage of each Zt469 unitig with the heterochromatin methylations H3K27me3 (orange bars) and H3K9me3 (purple bars), and the euchromatin methylation H3K4me2 (green bars) marks relative to unitig length. Unitig 9 is indicated by a black arrow. In both panels, except for unitig 9, only unitigs syntenic to the *Z. tritici* IPO323 reference genome are shown.

In line with these results, *in vitro* chromatin immunoprecipitation in combination with high-throughput sequencing (ChIP-seq) targeting three histone methylation marks (H3K4me2 for euchromatin and H3K27me3 and H3K9me3 for facultative and constitutive heterochromatin, respectively) also revealed that unitig 9, together with unitigs 18, 19 and 20, are enriched with the H3K27me3 mark, corroborating the low transcriptional activity observed for these genomic regions (Figure 3B). We also found the typical signatures of accessory chromosomes on unitig 17 (Figure 3A, B). Interestingly, this unitig is synthetic with the right chromosome arm of chromosome 7 in the *Z. tritici* IPO323 reference genome (Figure 1). It has been previously proposed that chromosome 7 in IPO323 may represent a fusion event of an accessory chromosome to a core chromosome (17), and our findings add further support to this hypothesis. Altogether, our results suggest that the novel unitig 9 along with unitigs 18, 19 and 20 represent accessory chromosomes in *Aegilops*-infecting *Z. tritici* isolates and carry similar hallmarks to the known accessory chromosomes in other *Zymoseptoria* species.

### High proportion of unitig 9 genes have orthologs in the sister species *Zymoseptoria ardabiliae*

Based on the previous finding that only few genes in unitig 9 have homologs in the reference *Z. tritici* IPO323 isolate, we further inspected the orthology of unitig 9 genes in other *Z. tritici* isolates and *Zymoseptoria* species. We used previously published and annotated whole genome assemblies based on long-read sequencing (PacBio) of the *Z. tritici* isolates Zt05 and Zt10 and of the closely related species *Zymoseptoria brevis* (Zb87), *Z. passerinii* (Zpa63), *Z. pseudotritici* (Zp13) and *Z. ardabiliae* (Za17) (29). Moreover, we included a new long-read, high quality genome assembly of the *Z. ardabiliae* isolate Za100 (Supplementary Table S1).

We identified 7,703 orthologous genes (i.e. orthogroups) distributed among all nine *Zymoseptoria* genome assemblies analyzed (Supplementary Figure S4A). Within species, we detected 146 species-specific orthogroups among *Z. tritici* isolates in comparison to 429 species-specific orthogroups shared between the two *Z. ardabiliae* isolates analyzed (Supplementary Figure S4A). Surprisingly, we found that a large number of orthogroups were specifically shared between Zt469 and Za100 (141 orthogroups), just a few orthogroups less than the number of orthogroups shared between all *Z. tritici* genomes (146 orthogroups) (Supplementary Figure S4A). In fact, comparisons between the *Z. tritici* isolates IPO323 and Zt469 and two *Z. ardabiliae* isolates (Za17 and Za100) revealed a considerably larger number of orthogroups shared between the *Aegilops*-infecting *Z. tritici* isolate Zt469 and the *Z. ardabiliae* isolates (number of shared orthogroups Za17: 59 and Za100: 240) as when compared to the intersection between IPO323 and the same *Z. ardabiliae* isolates (number of shared orthogroups; Za17: 37 and Za100: 82; Supplementary Figure S4B).

We next focused on the genes on unitig 9 in Zt469 and investigated their proportion in other *Zymoseptoria* genomes. As a comparison, we also analyzed the proportion of genes present in another accessory (unitig 19) and core chromosome (unitig 5) of Zt469 which are shared with the same *Zymoseptoria* isolates. As mentioned above, few orthogroups were observed between unitig 9 of Zt469 and the reference *Z. tritici* isolate IPO323 (only 4.3% orthogroups from unitig 9 were also present in the IPO323 genome) (Figure 1 and Figure 4). Similarly, we observed that the proportion of unitig 9 genes present in other *Z. tritici* genomes and in genomes of other closely related *Zymoseptoria* species was low, ranging from 1.7% in Zt10 (*Z. tritici*) to 4 % in Zpa63 (*Z. passerinii*) (Figure 4). Unexpectedly however, we observed a high proportion of unitig 9 genes with orthologs in the *Z. ardabiliae* isolates Za17 and Za100. Almost half of the genes in unitig 9 had orthologs in Za100 (47.2%), while Za17 had a lower, but still, compared to the other genomes, high proportion of orthologous genes (6.7%) (Figure 4). This high proportion of shared orthologs between unitig 9 of Zt469 and the Za100 isolate led us to the hypothesis that unitig 9 may have been exchanged between *Z. tritici* and *Z. ardabiliae* by introgression.

**Figure 4.**
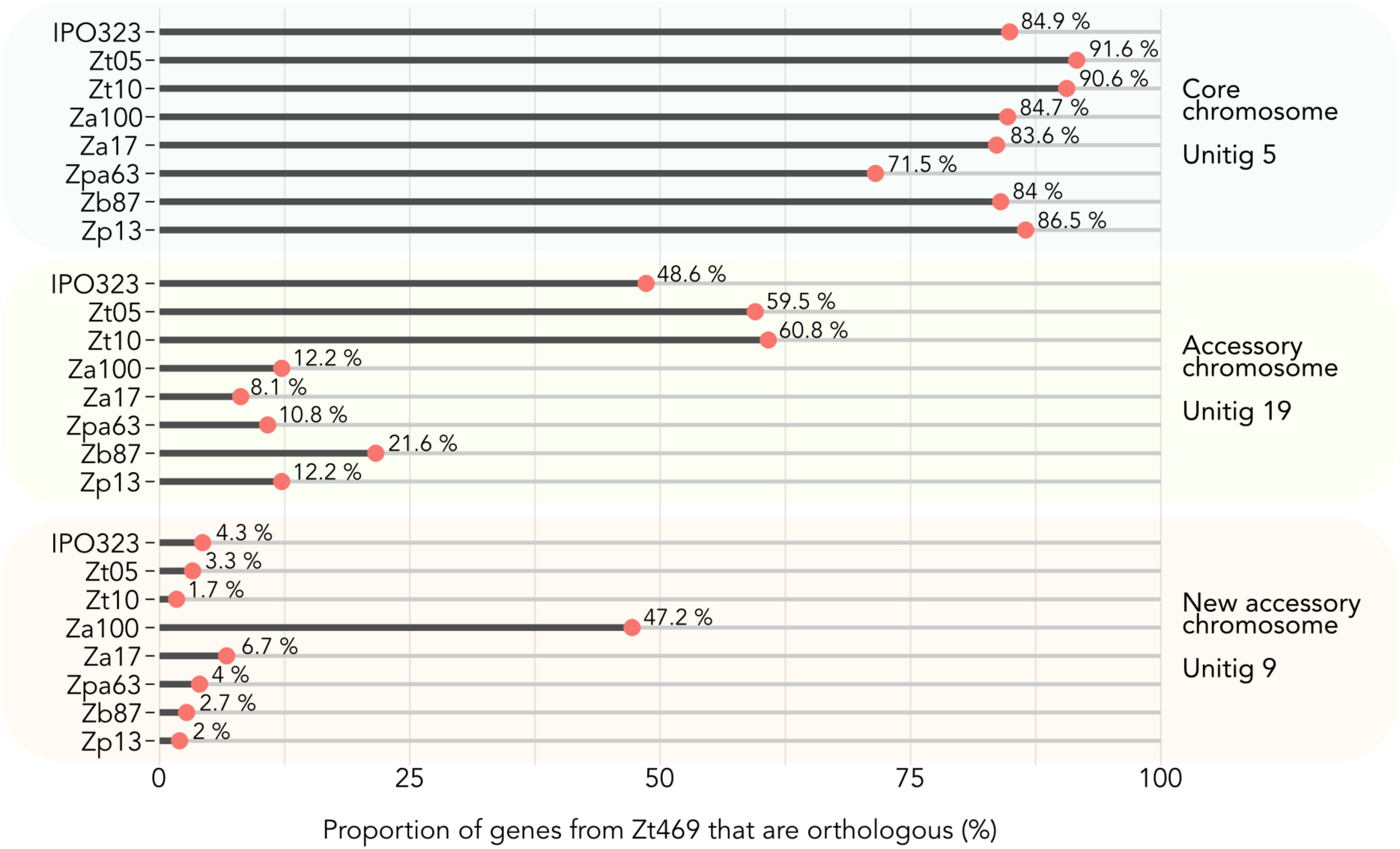
Proportion of Zt469 genes present in other *Zymoseptoria* species genomes. Proportion of genes present in unitig 5 (core), unitig 19 (accessory) and the newly described accessory unitig 9 of Zt469 and shared with other *Zymoseptoria* isolates was calculated for each genome individually. Only orthogroups solely shared between each unitig and the respective isolate genome are considered.

### An accessory chromosome of Za100 is syntenic with unitig 9 in Zt469

The observation that almost half of the proportion of genes in unitig 9 had orthologs in Za100 prompted us to further explore the distribution of orthologous genes in the *Z. ardabiliae* genome. As a first step, we analyzed chromosomal synteny between Za100 and Zt469 using the PacBio long-read assemblies and gene annotations. To this end, we plotted the synteny of orthologous genes between the two assemblies using a Circos plot (54). Contrasting to the comparison between the *Z. tritici* isolates Zt469 and IPO323 (Figure 1), we find that unitig 9 to a large extent is syntenic to unitig 3 in Za100, a presumably completely assembled chromosome comprising telomeric repeats in both ends (Figure 5). Intriguingly, the *Z. ardabiliae* chromosome is significantly larger than unitig 9 in length (∼2.6Mb compared to ∼1.6Mb). In spite of the size difference, we find that 95 % (134/141) of the genes of unitig 9 previously observed to have orthologs in Za100 (Figure 4) localize on unitig 3.

**Figure 5.**
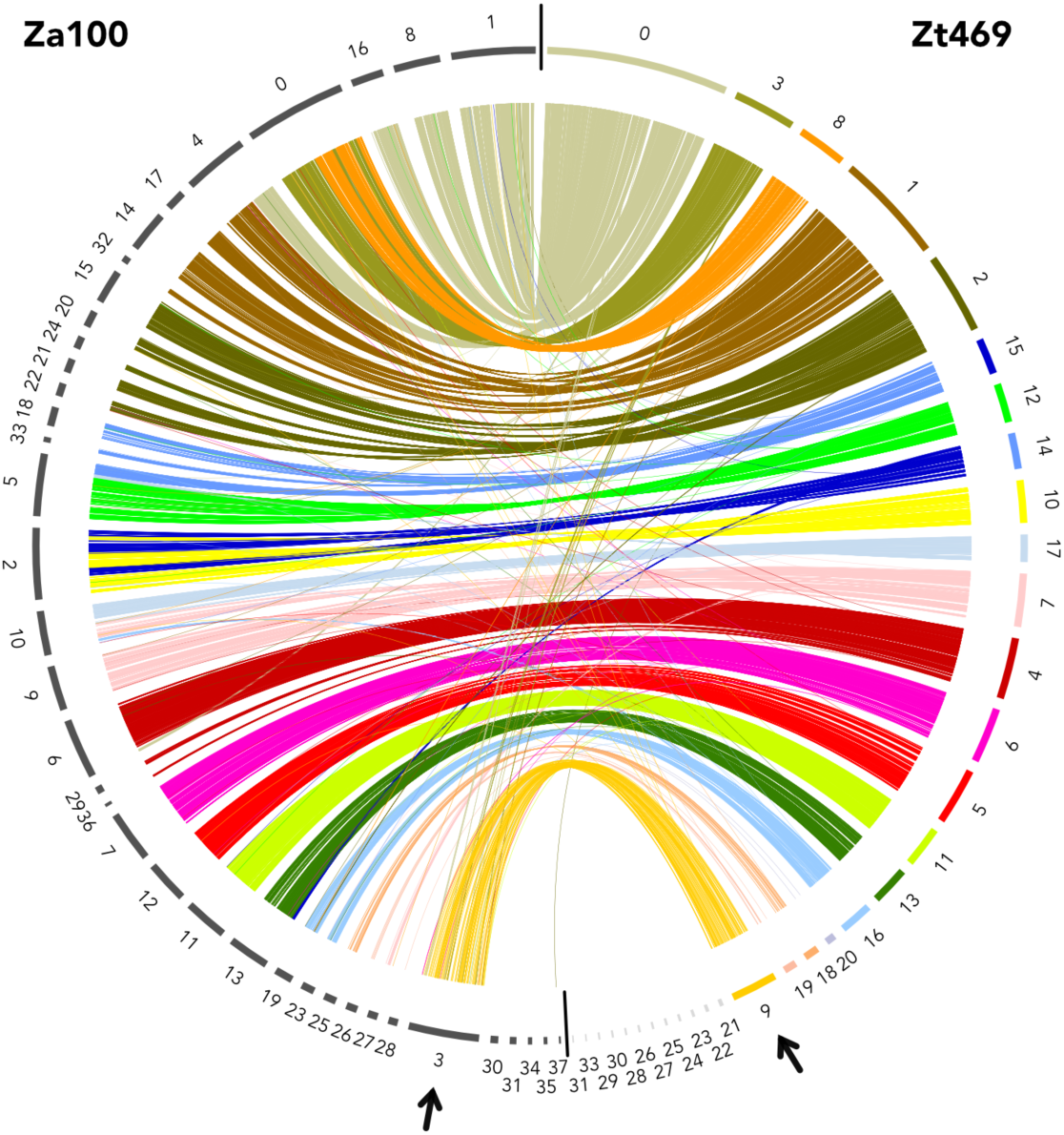
Unitig 3 in Za100 is syntenic to unitig 9 in Zt469. Synteny analysis was performed between the genomes of the *Aegilops*-infecting *Z. tritici* isolate Zt469 (right) and the *Z. ardabiliae* isolate Za100 (left). Each color represents a different contig in Zt469. Small Zt469 contigs are in light gray. Unitig 3 in Za100 shows synteny to unitig 9 in Zt469 (black arrows). Links represent orthologous genes. Contigs in Zt469 are ordered following the synteny to the reference IPO323 genome (Figure 1).

To further characterize the occurrence of unitig 3 in other *Z. ardabiliae* isolates, we determined the presence of unitig 3 in additionally 16 previously published *Z. ardabiliae* genomes collected across different years and locations in Iran (38, 41, 50). Using the PacBio Za100 assembly as reference and a normalized-read-coverage approach, we find that unitig 3 is present in only three additional isolates in our collection (Za19, Za20 and Za101) (Supplementary Figure S5). Similar results were observed when the same *Z. ardabiliae* isolates were mapped to the PacBio Zt469 genome assembly; hereby only these four isolates, as expected, showed the presence of a genomic fragment similar to unitig 9 as reflected by reads mapping to this unitig (Supplementary Figure S6). The PAV of unitig 3 among *Z. ardabiliae* isolates collected in different years (2004 and 2011) and locations led us to the conclusion that unitig 3 represents an accessory chromosome at low frequency in *Z. ardabiliae*.

### Unitig 3 in *Z. ardabiliae* has accessory chromosome hallmarks but is not enriched with the H3K27me3 methylation mark

In order to better characterize unitig 3 in Za100, we analyzed the genomic landscape of this chromosome. Similar to unitig 9 in Zt469, we observed a low gene density (average of 0.44 genes/kb), high TE density (average of 0.34 TEs/kb) and low transcriptional activity when compared to unitigs of core chromosomes and of similar size (e.g. unitig 4; 0.86 genes/kb and 0.20 TEs/kb on average) (Figure 6A). These signatures are consistent with other accessory chromosomes in *Zymoseptoria* species (28, 29, 31). In our search of accessory chromosome signatures, we also identified several other accessory unitigs represented by the shorter unitigs 24 to 37 with telomeric repeats in both ends in four of them (Figure 6A). Intriguingly, we observed that these accessory unitigs do not all comprise the same chromatin patterns, particularly unitig 3. ChIP-seq results targeting three histone methylation marks (H3K4me2, H3K27me3 and H3K9me3) revealed that unitig 3 is not enriched in the heterochromatin methylation mark H3K27me3 when compared to the other accessory unitigs (unitigs 25 to 37; Figure 6B). We also observed accessory chromosome signatures on unitig 10, which is syntenic to unitig 17 in Zt469 (Figure 5), and to the “accessory arm” of chromosome 7 in *Z. tritici* IPO323 (Figure 1; (17). The low enrichment of H3K27me3 in unitig 3 suggests that, even though the unitig has some accessory chromosome-hallmarks (high TE, low gene density and low transcriptional activity), it still has a different profile of histone modifications compared to other accessory chromosomes described in *Zymoseptoria* species.

**Figure 6.**
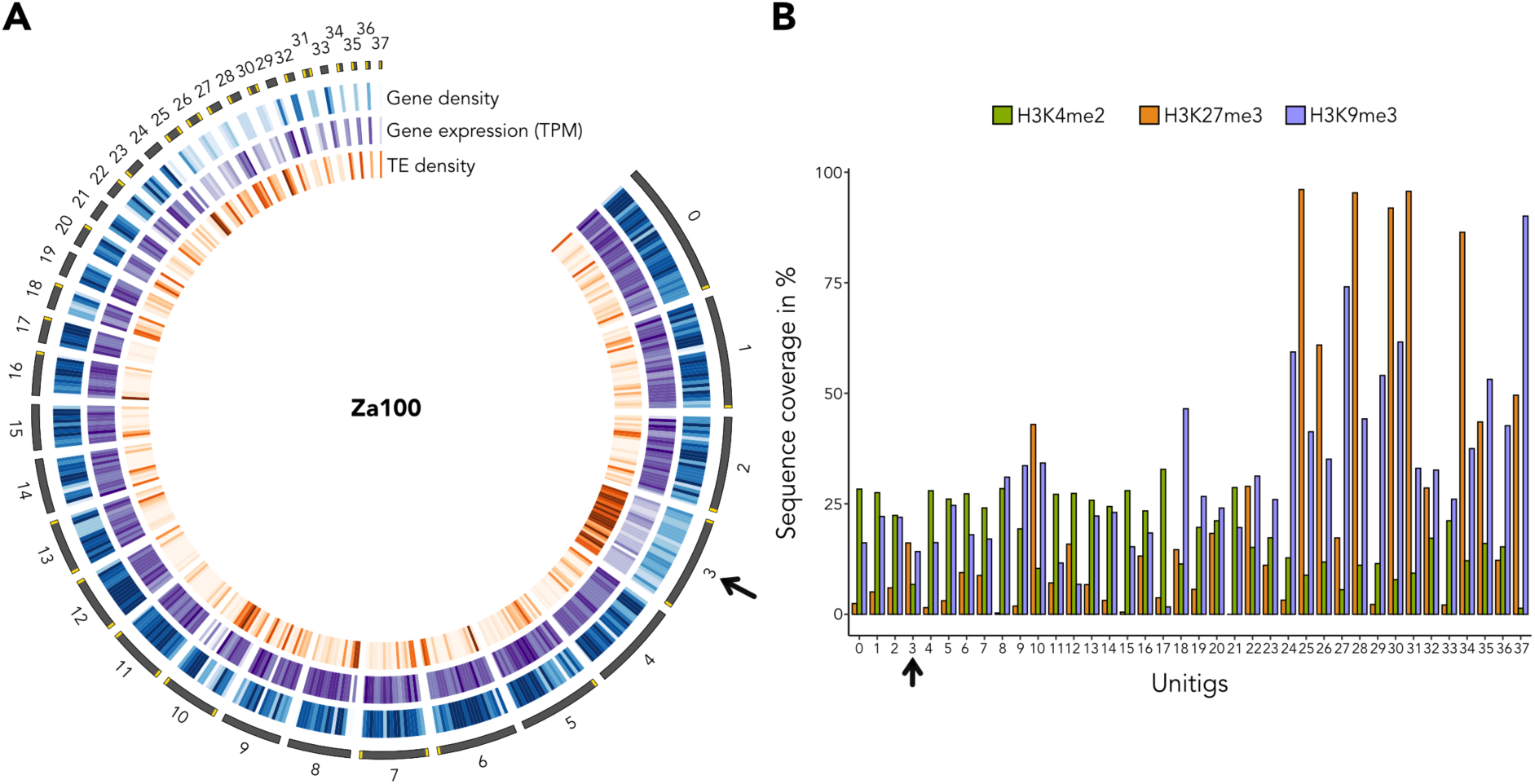
Unitig 3 shows intermediate accessory chromosome hallmarks. **(A)** Circos plot representing different features along the thirty-eight Za100 unitigs. Unitigs are represented by the dark gray segments sorted by length. Telomeric repeats are indicated in yellow. Tracks from outside to the inside are heatmaps in 100kb windows representing respectively: gene density, gene expression *in vitro* in log2(TPM+1) and TE density. Darker heatmap colors indicate higher values. Unitig 3 is indicated by a black arrow. **(B)** Barplot displaying the percentage of sequence coverage of each Za100 unitig with the heterochromatin methylations H3K27me3 (orange bars) and H3K9me3 (purple bars), and the euchromatin methylation H3K4me2 (green bars) marks relative to unitig length. Unitig 3 is indicated by a black arrow.

### Unitig 3 and unitig 9 show high transcription activity of TEs and different genome defense signatures

Considering the differences observed in length and histone methylations marks between unitig 3 in Za100 and unitig 9 in Zt469, we focused on further characterizing the TE landscape of these genomes. TEs can promote intra- and inter-specific variability in terms of genome structure, size and transcriptional regulation, and their proliferation control can be tightly linked with DNA methylation or heterochromatin-associated histone modifications (12, 99–101).

Analyses of TE content in Zt469 and Za100 revealed that the two genomes vary in the overall proportion of TEs; 19,02% in Za100 compared to 15,76% in Zt469 (Supplementary Figure S7A). Comparing the composition of TE families in the two genomes, we find, in agreement, with previous studies a high proportion of LTR-retrotransposons compared to other TE families (Supplementary Figure S7B) (45). Regarding unitig 3 and unitig 9, although both accessory unitigs are enriched in TEs (Figures 3 and 6), unitig 3 in Za100 shows a higher TE proportion (15,36% of TEs and approximately 0.39 Mb of the chromosome length) compared to unitig 9 in Zt469 (13.36% of TEs and approximately 0.2 Mb of the chromosome length) (Supplementary Figure S8A). In unitig 3, retrotransposons from the LINE order make up the largest portion of TEs while for unitig 9 most of the TEs could not be categorized (“noCat”; Supplementary Figure S8B) suggesting the activity of different elements on the two homologous chromosomes in the two fungal species.

Interestingly, not only do the unitigs differ in TE proportion but they also differ in the levels of expression of these genomic elements during *in vitro* growth. Although both unitigs show a high TE expression, a comparative analysis of TE expression based on RNA-seq data showed significantly higher levels of expression of TEs on unitig 3 in *Z. ardabiliae* compared to expression of TEs on unitig 9 in *Z. tritici* (Wilcoxon rank sum test, p-value=0.0016) (Supplementary Figure S9). For both unitig 3 and unitig 9, expression levels of TEs were in general significantly higher than expression levels of genes within the same chromosome (Wilcoxon rank sum test, p-value < 2.22e-16) (Supplementary Figure S9B). Based on these results, we hypothesized that unitig 3 and unitig 9 have a reduced efficacy of genome defense mechanisms regulating TE activity.

To further test this hypothesis, we assessed signatures of another genome defense mechanism known to be prominent in many ascomycetes, namely Repeat Induced Point (RIP) mutations (102, 103). To quantify putative RIP-derived mutations, we scanned each genome individually and searched for dinucleotide bias mutations in duplicated sequences. Based on this, we calculated the RIP compositive indices and determined the distribution of large RIP affected regions (LRARs) in 1 kbp genomic windows using the RIPper software (89). At genome-wide level, we observed that RIP signatures were present in 21.85 % of the total genomic content of Za100 and 18.61 % of the total genomic content in Zt469 (Supplementary Table S7). Large RIP affected regions (LRARs), besides comprising extensive AT-rich regions, have an average size of 23585 bps and 18299 bps and affected a total of 8.2 Mbp and 6.9 Mbp in the genomes of Za100 and Zt469, respectively (Supplementary Table S7). Intriguingly, the proportion of RIP mutations differed considerably between the two species for the two accessory unitigs 3 and 9: at the chromosome level, the overall percentage of 1 kbp regions per unitig affected by RIP ranged from 2.72% to 85.52% in Za100 and from 7.22% to 31.5% in Zt469 (Supplementary Figure S10). RIP signatures specifically in unitig 3 and unitig 9 comprised 24.09% and 28.95% of the total unitig contents, respectively (Supplementary Figure S10).

We also estimated RIP composite indices per 50bp windows of each TE copy in the genomes of both Zt469 and Za100 and observed that, on average, TE copies were statistically more RIPped in unitig 9 of Zt469 compared to TEs in unitig 3 of Za100 (Wilcoxon rank sum test, p-value < 2.22e-16; Figure 7). The same pattern was observed when including genome-wide TE copies (Figure 7). Altogether, our results suggest that the efficacy of RIP has been higher for TEs on unitig 9 compared to unitig 3. The lower extent of H3K27me3 and the overall higher transcriptional activity of TEs in unitig 3 (Supplementary Figure S9) corroborates our deduction that unitig 3 of Za100 is enriched with active TEs.

**Figure 7.**
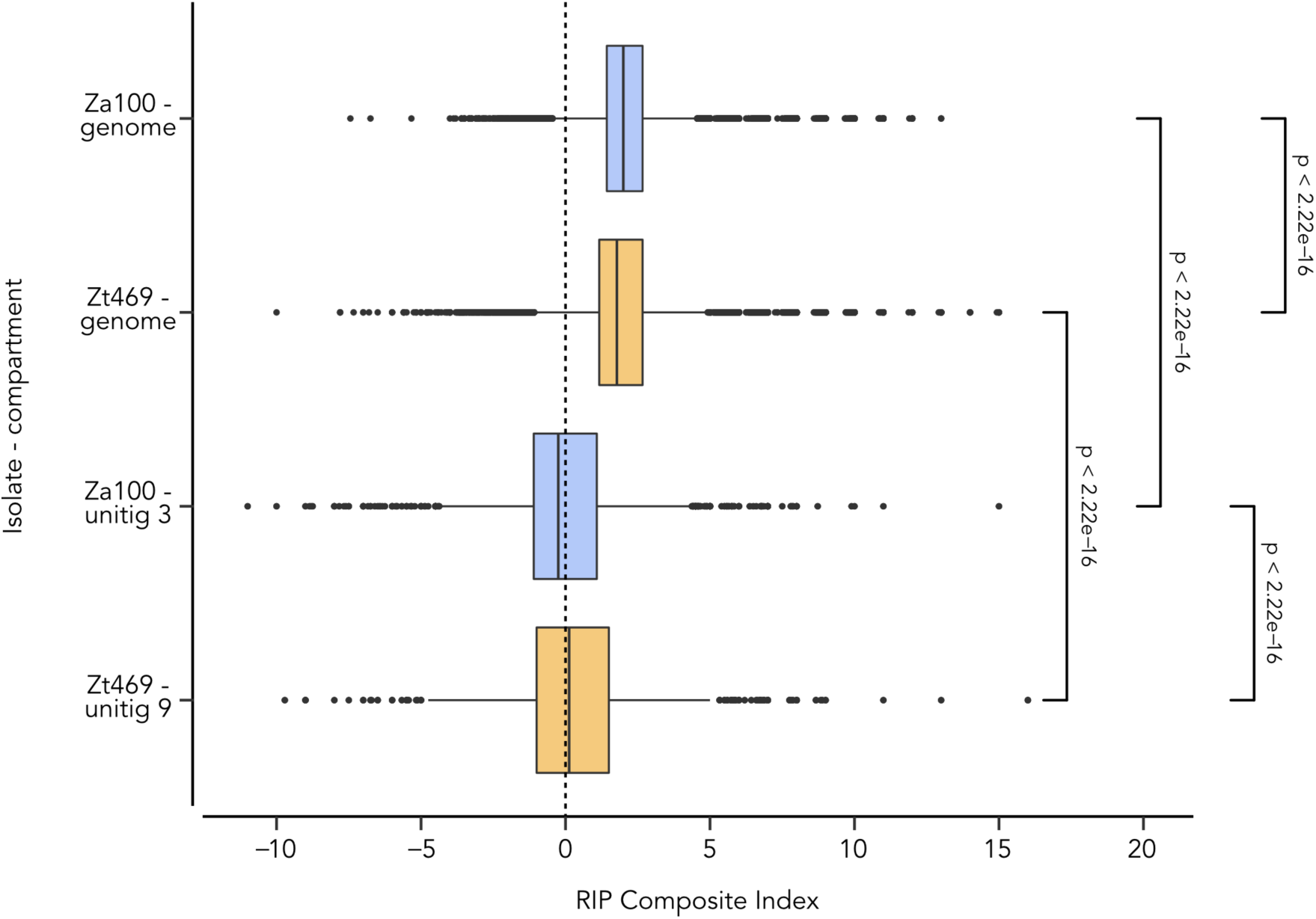
Repeat-Induced Point (RIP) mutation signatures in TE copies of Za100 and Zt469 genomes. Distribution of RIP composite indices calculated in 50bp sliding windows per TE copy distributed genome-wide (top two boxes) and per TE copies present on the unitigs 3 and 9 (bottom two boxes) for *Z. ardabiliae* Za100 (blue color) and *Z. tritici* Zt469 isolates (orange color). Vertical dashed line represents the threshold (0) above which composite indices values indicate RIP signature. *P*-values were calculated using pairwise Wilcoxon rank sum tests.

### Gene expression in unitig 9 suggests a possible role during infection

We finally asked if unitig 9 in *Z. tritici* Zt469 has a functional relevance by analyzing gene and TE expression patterns during *in vitro* growth and during *in planta* infection. For expression profiling *in planta*, we analyzed stage-specific RNA-seq datasets based on four infection stages based on previously generated transcriptome data (46). To determine the levels of expression in individual genes and TEs, we used the gene and TE models described previously and functionally annotated each gene using different tools (see Methods and Supplementary Tables S2, S3 and S5). Expression levels, based on three biological replicates per condition, were compared using normalized read mappings to transcript per million (TPM) in log2(TPM+1). The percentage of reads that could be mapped to the Zt469 genome were on average 93.4 % between replicates for *in vitro* growth and ranged from 2.3 % to 76.5 % between replicates and infection stages during *in planta* infection (Supplementary Table S8).

At first, we focused on the overall expression levels of genes and TEs in unitig 9 and asked if the expression of these features was different between *in vitro* and *in planta* conditions. We observed a lower transcriptional activity of genes compared to TEs not only *in vitro*, but also *in planta* (Figure 3; Supplementary Figure 11 and Supplementary Table S9). Previous transcriptome studies of *Z. tritici* during *in vitro* growth and *in planta* infection have shown that, on average, the expression levels of genes located on accessory chromosomes were up to 20-fold lower (in RPKM) than the expression levels of genes present on core chromosomes (31, 32). Our results corroborate these findings by demonstrating that genes located in unitig 9, and also in the other accessory unitigs 18, 19 and 20, were expressed on average from 1.6- to 2.1-fold lower levels (in TPM) than the genes located in the core chromosomes (unitigs 0 to 17; Supplementary Tables S9 and S10).

Next, we functionally annotated the gene models in the Zt469 genome and focused on the expression profile of specific gene categories in unitig 9 (see Methods and Supplementary Tables S2, S3 and S5). We identified 26 genes with putative functions in host-pathogen interactions including secreted proteins as CAZymes (carbohydrate-degrading enzymes), effectors and other small secreted proteins (SSPs) as well as secondary metabolites encoded by biosynthetic gene clusters (BGCs) (102–105). Of the 26 genes (Supplementary Table S5), 16 genes showed no transcription (TPM=0) in at least one condition (Figure 8). Most of the 26 genes analyzed are not transcribed at infection stage “A” (7dpi) followed by a low to intermediate expression (0.03 < log2(TPM+1) < 0.030) at infection stage “D” (21dpi; Figure 8). Taking these results together, we find that genes on unitig 9 show differential expression profiles during disease progression suggesting a possible role in plant infection.

**Figure 8.**
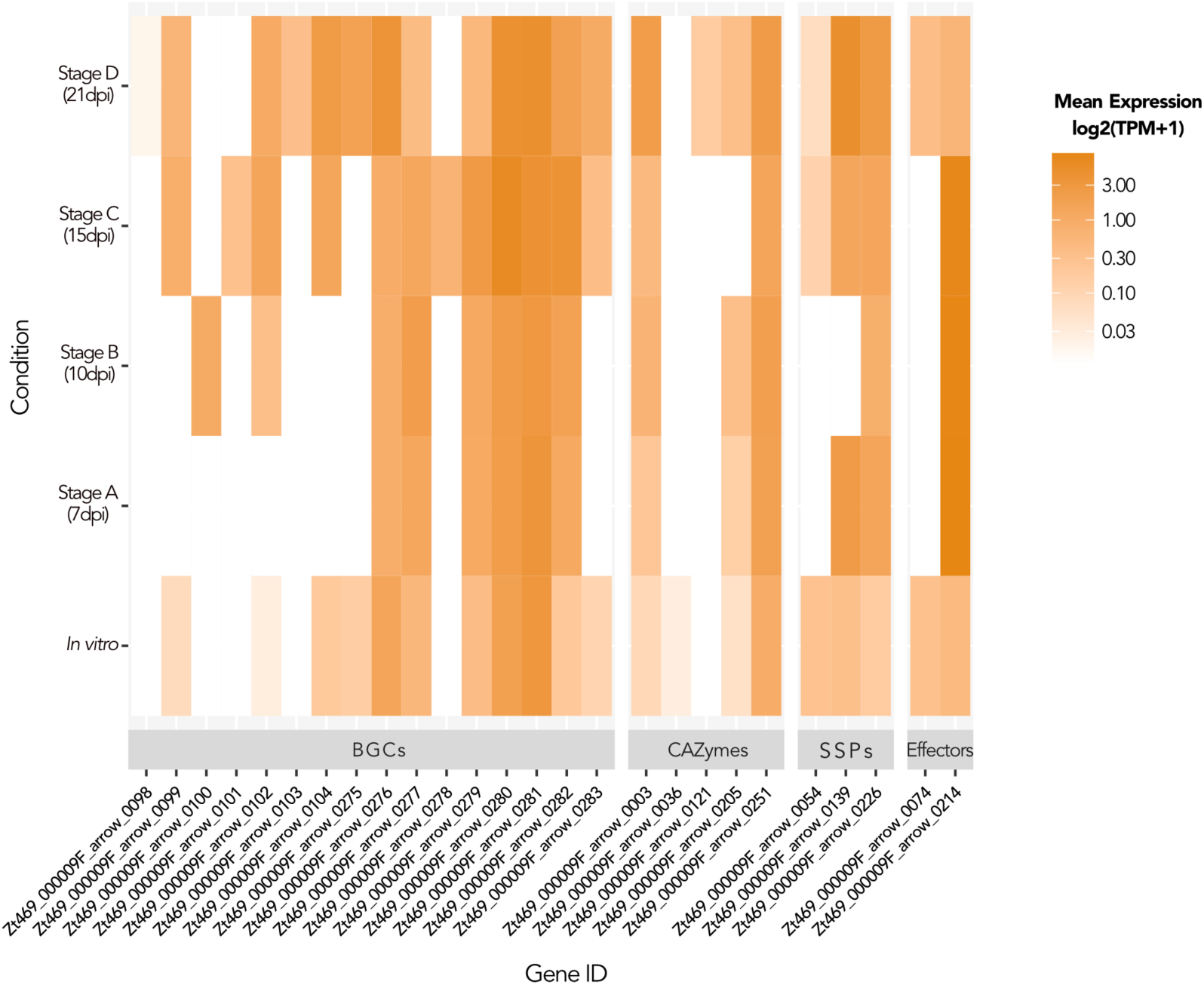
Gene categories involved in host-pathogen interactions show different levels of expression in unitig 9 during *in vitro* growth and *in planta* infection. Heatmaps showing mean expression levels in log2(TPM+1) for each gene individually in unitig 9 belonging to four gene categories analyzed at different infection stages *in planta* and *in vitro*. White cells represent no transcription (TPM=0). BGCs (Biosynthetic Gene Clusters); CAZymes (Carbohydrate-degrading enzymes); SSPs (Small Secreted Proteins).

### Introgression tests suggest gene flow between *Aegilops*-infecting *Z. tritici* and *Z. ardabiliae*

Considering the synteny between the unitig 9 of Zt469 and unitig 3 in Za100, we hypothesize that these accessory chromosomes may have been exchanged by introgression between *Aegilops*-infecting *Z tritici* and *Z. ardabiliae* isolates. To test this hypothesis, we performed an ABBA-BABA test (94, 95), also known as Patterson’s *D*-score, among four taxa (P1, P2, P3, O) to determine whether interspecific gene flow occurs. To this end, we included the *Aegilops*-infecting *Z. tritici* isolates Zt469 (with unitig 9) or Zt436 (without unitig 9) as P1; the wheat-infecting *Z. tritici* isolates Zt565 or Zt668 as P2; and the *Z. ardabiliae* isolates Za100 (with unitig 3) or Za98 (without unitig 3) as P3 (Supplementary Figure S1). We included *Aegilops*-infecting Z*. tritici* isolates with or without unitig 9 as well as *Z. ardabiliae* isolates with or without unitig 3 to evaluate whether gene flow only have occurred between isolates harboring these newly described chromosomes or if signatures of introgression could be detected generally at the lineage and species levels. All tests were performed including the genome of the *Z. passerini* isolate Zpa796 as the outgroup (O).

Using the tree topologies, we observed significantly negative *D* statistic results (Z-score > |3|) for tests including *Aegilops*-infecting *Z. tritici* isolates with or without unitig 9, which indicates an excess of shared derived alleles between these Z*. tritici* isolates and *Z. ardabiliae* (Supplementary Table S11). We also observed significantly negative *D* statistic results (Z-score > |3|) if *Z. ardabiliae* isolates with or without unitig 3 were included in the tests (Supplementary Table S11). These results were consistent when we replaced Zt565 with another wheat-infecting *Z. tritici* isolate, Zt668 (Supplementary Table S11). These findings suggest that interspecies hybridization occurs between *Aegilops*-infecting *Z. tritici* and *Z. ardabiliae* species and it is not only restricted to isolates harbouring the newly described chromosomes.

At last, we tried to access the relative age and possible introgression direction of the syntenic chromosomes in *Aegilops*-infecting *Z. tritici* and *Z. ardabiliae*. We calculated the levels of genomic variation in each chromosome using population data, and used nucleotide diversity (*π*) per 1 kb windows as a proxy of age. Hereby, we hypothesized that an older chromosome would accumulate more mutations and therefore contain higher levels of nucleotide diversity as compared to a younger chromosome. Using quality-filtered SNPs, we observed a statistically higher nucleotide diversity in unitig 9 of Zt469 than unitig 3 in Za100 (Wilcoxon rank sum test, p-value < 0.001; Supplementary Figure S12). Based on our hypothesis, these results suggest that unitig 9 accumulated more polymorphisms and therefore may be older than unitig 3. This would imply that the chromosome originated first in *Z. tritici* and subsequently was transferred to *Z. ardabiliae* by introgression.

## Discussion

Accessory chromosomes frequently found in fungi provide a source of genetic innovations considering the low density of “essential” genes, and therefore a reduced efficacy of purifying and background selection. Moreover, the activity of transposable elements (TEs) can create new genetic variants (1, 14, 15). In this study, we characterize a new and unique accessory chromosome in *Aegilops*-infecting *Z. tritici* and in the closely related species *Z. ardabiliae* using a multi-omics approach and high-quality whole genome assemblies, gene and TE annotations. Our results confirm that recurrent interspecies hybridization occurs between *Zymoseptoria* species in natural vegetations and may serve as a route for accessory chromosomes emergence.

Consistent with previous analyses on the genomic structure of *Zymoseptoria* species (17, 29, 31, 32, 40), the genome architecture in the analyzed *Zymoseptoria* isolates is also highly compartmentalized. We demonstrate that unitig 9 and the smaller unitigs 18, 19 and 20 in the *Aegilops*-infecting *Z. tritici* isolate Zt469 exhibit accessory chromosome hallmarks as described previously in the wheat-infecting *Z. tritici* and closely related *Zymoseptoria* species, including enrichment in heterochromatin methylation mark H3K27me3; PAV between isolates and low levels of gene transcription *in vitro* and *in planta* (17, 29, 32). Similarly in the *Z. ardabiliae* Za100 isolate, accessory unitigs, including the large unitig 3, showed characteristics that distinguish them from the core unitigs as e.g. PAV between isolates; low gene and high transposon densities; and low transcriptional gene activity. Considering the divergence observed between the *Aegilops*- and wheat-infecting *Z. tritici* isolates (46) and between the *Z. tritici* and *Z. ardabiliae* species (4, 25, 38, 39), these results corroborate previous findings that genome compartmentalization and accessory chromosomes are ancestral traits among *Zymoseptoria* species (29).

We observed interesting patterns regarding TE suppression on unitig 3 of Za100. One of the crucial roles of heterochromatin in eukaryotes is to prevent genome instability by the silencing of TE replication and spread (104). In several fungal plant pathogens, including *Zymoseptoria* species, enrichment of the facultative heterochromatin methylation mark H3K27me3 has been considered a hallmark of accessory chromosomes and can explain the overall transcriptional gene silencing observed for these chromosomes (13, 17, 105). H3K9me2/3 on the other hand has been demonstrated to be enriched in repeat-rich regions with a direct impact on transposable elements (TEs) control and genome stability (106–109). Transcriptional activation of TEs can result in their spread throughout the host genome, causing insertional mutations and chromosomal rearrangements (110–112). Moreover, several ascomycete fungi comprise a unique genome defense mechanism, known as RIP (Repeat-Induced Point mutations), to mutate and thereby inactivate repetitive sequences (102, 103). Interestingly, although we observed that unitig 3 shows most of the expected features of an accessory chromosome, this unitig was not enriched with the heterochromatin methylation marks H3K27me3 and H3K9me3. We also observed that unitig 3 has reduced RIP signatures when compared to most small and accessory unitigs in Za100 and to the syntenic unitig 9 in Zt469. In line with these results, we found that the levels of TE transcription in unitig 3 overall are high and even superior to the transcription of genes on the same unitig and to the levels observed for unitig 9 in Zt469. Taking these results together, we conclude that unitig 3 lack efficient silencing mechanisms, which may have resulted in the spread of TEs along the unitig and rapid sequence diversification. The relevance of distinct genome compartments in the maintenance of active TEs has also been demonstrated in flies. In male *Drosophila* flies, it was found that the “toxic” repeat-rich Y chromosomes act as reservoirs of actively transcribed TEs that may lead to reduced male fitness by deleterious TE mobilization and ineffective heterochromatin silencing of these repetitive elements (113, 114). Further analyses on Za100 TEs activity and their genomic and epigenetic control *in vitro* and potentially *in planta* can shed a light on the dynamic of TEs and their impact on *Z. ardabiliae* development and fitness.

Exchange of chromosomes between pathogen lineages or species through introgression can generate novel pathotypes and ultimately new pathogen species (115–117). Experimental evidence from *Fusarium oxysporum* demonstrates that accessory (or lineage-specific) chromosomes can be vertically transferred between distinct lineages by hyphal fusion and convert non-pathogenic strains into a pathogen in specific hosts (3, 118). Other reports have also shown that horizontal transfer of chromosomes between fungal strains can occur in *Colletotrichum gloeosporioides* (119), in *Alternaria alternata* (120) and in the cereal blast pathogen *Magnaporthe oryzae* (121). In *Z. tritici*, it has been suggested that accessory chromosomes have originated via ancient horizontal transfer from an unknown donor followed by extensive recombination events (28). However, other studies have pointed to alternative mechanisms. One study demonstrated that a *Z. tritici* accessory chromosome emerged by non-disjunction duplication of core chromosomes followed by a degeneration process via breakage-fusion-bridge (BFB) cycles and RIP on duplicated sequences (21). Here we demonstrate the introgression of chromosomes between *Zymoseptoria* species followed by TE activation may be an alternative mechanism whereby new accessory chromosomes can arise and diversify.

The relevance of introgression in accessory chromosome evolution is apparent from the comparison of genome sequences across diverse *Zymoseptoria* genomes. First of all, we found that the accessory unitig 9 in *Aegilops*-infecting *Z. tritici* is syntenic to another accessory chromosome (unitig 3) in the *Z. ardabiliae* species. The lack of further population data of *Z. ardabiliae* isolates harbouring unitig 3 prevented us to explore whether the chromosome itself represents an introgressed genomic segment as well to determine the time and direction of introgression. However, we found evidence that genome-wide introgression signatures are present between *Aegilops*-infecting *Z. tritici* and *Z. ardabiliae* species through ABBA-BABA tests, which suggests that interspecies hybridization takes place and the transfer of chromosomes are likely to occur between these two species, particularly when considering their sympatry in grassland vegetations in the Middle East (39, 46).

Considering the findings we gathered so far, some hypothetical introgression scenarios can be made about the evolutionary origin of the novel accessory chromosomes in *Z. tritici* and *Z. ardabiliae* species:

In a first scenario, we consider that these newly described chromosomes were acquired independently from a common ancestral *Zymoseptoria* population and since then have been segregating only in *Aegilops*-infecting *Z. tritici* and *Z. ardabiliae* populations. This scenario could be supported by the host range and evolutionary history of the *Zymoseptoria* genus in the Middle East. It is possible that the newly described unitig 9 and unitig 3 persisted in wild grass-infecting populations of *Z. tritici* and *Z. ardabiliae* from the last common ancestral *Zymoseptoria* population that also infected wild grasses. This scenario is however incompatible with the unexpected sequence similarity of unitig 3 and unitig 9.

In a second scenario, chromosomal introgression could have happened from *Z. ardabiliae* into *Aegilops*-infecting *Z. tritici*. In this scenario, we may assume that unitig 3 is a chromosome that remained from ancestral populations and it is being lost in the *Z. ardabiliae* species, resulting in the low frequency of the unitig among *Z. ardabiliae* isolates in our collection. The detrimental effects of active TEs (110–112) and the lower efficacy of genome defenses against TE replication (e.g. heterochromatin-associated methylation and RIP) on unitig 3 may have led to selection against this chromosome over *Z. ardabiliae* generations. The considerably lower proportion of TEs on the introgressed chromosome however argues against this scenario.

In a third introgression scenario, we consider the possibility that a recent chromosomal introgression happened from *Aegilops*-infecting *Z. tritici* isolates into the *Z. ardabiliae* species. This scenario can be supported by the low frequency of unitig 3 observed among our collection of *Z. ardabiliae* isolates spanning different years and locations, suggesting that this chromosome has not been established yet at the *Z. ardabiliae* species level. The higher degree of nucleotide diversity observed in unitig 9 compared to unitig 3 also suggests that unitig 9 in *Aegilops*-infecting *Z. tritici* isolates accumulated more polymorphisms and it is therefore potentially older than unitig 3 in *Z. ardabiliae*. Moreover, we also observed high TE transcription activity, low enrichment of the heterochromatin-associated methylation marks (H3K27me3 and H3K9me3) and low RIP signatures in unitig 3 and in the TE copies that reside within this chromosome, which could further corroborate that unitig 3 has been recently introgressed and since then acting as a reservoir of transcriptionally active TEs.

In light of our findings, we suggest that the third introgression scenario is the most parsimonious explanation for the origin and maintenance of the syntenic accessory chromosomes unitig 9 and unitig 3 in *Z. tritici* and *Z. ardabiliae* species. Further sampling of *Z. ardabiliae* isolates containing unitig 3 is required to better characterize the evolution of the newly acquired chromosome in this species.

## Conclusions

In this study we identified a unique, and so far, undescribed accessory chromosome in *Z. tritici* and *Z. ardabiliae* using a multi-omics comparative framework. Our findings provide valuable resources to investigate the evolutionary mechanisms and potential impact of interspecies hybridization and TE activity in the “birth” of new accessory chromosomes in these ascomycete fungi. The occurrence of these chromosomes only in fungal species infecting wild grasses highlights the importance of wild pathosystems on adaptive genome evolution of fungal pathogens and can further illustrate a possible source of genetic novelty for closely related fungal lineages infecting important crops like *Z. tritici* on wheat.

## Supporting information

Fagundes_2024_Zymo_NewChr_Supplementary_Material

## Acknowledgements

The authors would like to thank Cécile Lorrain for support with the TE analyses and the members of the Environmental Genomics group for helpful discussions. This research was funded by the Max Planck Society, Germany by a fellowship to EHS.

## Data availability

All short-read genomic sequences analyzed during this study are publicly available in different repositories. For accession numbers, repositories and references please see Supplementary Table S12. The Zt469 and Za100 assembled genomes can be found at https://doi.org/10.5281/zenodo.13773246. Gene and TE annotations for both Zt469 and Za100 are deposited at https://doi.org/10.5281/zenodo.13773246. *In planta* RNA-seq datasets for *Z. tritici* Zt469 from (46) are available at the NCBI SRA BioProject accession number PRJNA1162778. *In vitro* RNA-seq datasets for Zt469 and Za100 are available at the accession number NCBI SRA BioProject PRJNA1162778. The ChIP-seq datasets for both Za100 and Zt469 are available under the NCBI SRA BioProject accession number PRJNA1162778. The genome sequence of the reference *Z. tritici* isolate IPO323 (MYCGR v2.0) is available at NCBI under the RefSeq assembly GCF_000219625.1.

## Supplementary Material

### Supplementary Figures

**Figure S1. Tree topologies used in ABBA-BABA tests**. ABBA-BABA tests were performed among four taxa (P1, P2, P3, O) in eight tree topologies. We included the *Aegilops*-infecting *Z. tritici* isolates Zt469 (with unitig 9) or Zt436 (without unitig 9) as P1; the wheat-infecting *Z. tritici* isolates Zt565 or Zt668 as P2; and the *Z. ardabiliae* isolates Za100 (with unitig 3) or Za98 (without unitig 3) as P3. All tests included the genome of the *Z. passerini* isolate Zpa796 as outgroup (O).

**Figure S2. Presence of unitig 9 in Zt469 is confirmed by PFGE and Southern blot analyses.** To validate the presence of unitig 9 at the expected assembly size in Zt469, we performed Pulsed-Field Gel Electrophoresis (PFGE) **(A)** followed by Southern blot analyses **(B)** in non-protoplast plugs of this isolate and in the reference *Z. tritici* isolate IPO323 (strain Zt244); in the wheat-infecting *Z. tritici* isolate Zt10 (strain Zt366); and in the *Aegilops*-infecting *Z. tritici* isolate Zt501. A southern blot probe of 1.5 kb was designed to hybridize specifically in unitig 9 of Zt469 (Supplementary Table S6). Probe hybridization was detected at the expected unitig 9 size (∼1.6Mb; **B**). Chromosomal DNA of *Hansenula wingei* (Bio-Rad, Munich, Germany) was used as a standard size marker for PFGE.

**Figure S3. *Aegilops*-infecting *Z. tritici* isolates show high PAV of accessory chromosomes**. Chromosome presence-absence variation (PAV) in host-diverging *Z. tritici* populations was analyzed by read mapping to the *Z. tritici* IPO323 reference genome. Heatmap represents normalized mean coverage of reads mapped to each position of the reference genome. Darker colors represent regions of higher coverage as e.g. repetitive elements or duplications.

**Figure S4. A large number of orthologs is shared between Zt469 and Za100 isolates. (A)** Upset plot showing the number of orthogroups shared between the nine *Zymoseptoria* species genomes analyzed. Only intersects with more than 100 orthogroups are displayed. Number of orthogroups shared between Zt469 and Za100 are highlighted in red. **(B)** Venn diagram summarizing the orthogroups exclusively shared between the *Z. tritici* isolates IPO323 and Zt469 and *Z. ardabiliae* isolates Za17 and Za100.

**Figure S5. Unitig 3 shows presence-absence variation in *Z. ardabiliae* isolates**. Chromosome presence-absence variation (PAV) was analyzed by read mapping to the *Z. ardabiliae* Za100 PacBio genome assembly. Heatmap represents normalized mean coverage of reads mapped to each position of the genome assembly. Only unitigs larger than 100 kb are displayed and sorted by descending order of length. Darker colors represent regions of higher coverage as e.g. repetitive elements or duplications.

**Figure S6. Read mapping to Zt469 genome confirms synteny and PAV of unitig 9 among *Z. ardabiliae* isolates**. Chromosome presence-absence variation (PAV) was analyzed by read mapping to the *Z. tritici* Zt469 PacBio genome assembly. Heatmap represents normalized mean coverage of reads mapped to each position of the genome assembly. Except for unitig 9, only unitigs syntenic to the *Z. tritici* IPO323 reference genome are shown and sorted in descending order of length. Darker colors represent regions of higher coverage as e.g. repetitive elements or duplications.

**Figure S7. Transposable element (TE) content in Za100 and Zt469 genomes**. **(A)** Total percentage of genome covered by TEs in Za100 (left) and Zt469 (right). **(B)** Stacked bar plot showing the TE content (%) per genome. Colors represent TE order coverage with retrotransposons (LTR, LINE, TRIM, LARD and other class I orders) and DNA transposons (TIR, MITE, Helitron and Maverick). TEs that could not have been classified are indicated as “no category” (“noCat”).

**Figure S8. Transposable element (TE) content in the syntenic unitig 3 in Za100 and unitig 9 in Zt469**. **(A)** Total percentage of unitig covered by TEs in unitig 3 (left) and unitig 9 (right). **(B)** Stacked bar plot showing the TE content (%) per unitig. Colors represent TE order coverage with retrotransposons (LTR, LINE, TRIM, LARD and other class I orders) and DNA transposons (TIR, MITE, Helitron and Maverick). TEs that could not have been classified are indicated as “no category” (“noCat”).

**Figure S9. Unitig 3 shows higher transcriptional activity of TEs than unitig 9. (A)** Circos plots representing gene and TE expression *in vitro* in log2(TPM+1) along the Za100 (left) and Zt469 (right) genomes. Unitigs are represented by the dark gray segments sorted by length. For Zt469, except for unitig 9, only unitigs syntenic to the *Z. tritici* IPO323 reference genome are shown. Tracks from outside to the inside are heatmaps in 100kb windows representing respectively: gene expression *in vitro* in log2(TPM+1) and TE expression *in vitro* in log2(TPM+1). Darker heatmap colors indicate higher values. **(B)** Violin plots representing the mean expression levels in log2(TPM+1) for both gene and TE features of unitig 3 in Za100 (violet color) and unitig 9 in Zt469 (orange color) during *in vitro* growth. P-values were calculated using pairwise Wilcoxon rank sum tests.

**Figure S10. Unitig 3 and unitig 9 are differentially affected by RIP.** Lolliplot plots showing the percentage of regions (1 kbp windows) per unitig affected by RIP in Za100 **(A)** and Zt469 **(B)** genomes. Unitigs are sorted by descending order of length. For Zt469 **(B)**, except for unitig 9, only unitigs syntenic to *Z. tritici* IPO323 reference genome are shown.

**Figure S11. Genes and TEs show different levels of expression in Zt469 unitig 9 during *in vitro* growth and *in planta* infection.** Boxplots representing the mean expression levels in log2(TPM+1) for genes and TEs in unitig 9 at different infection stages *in planta* and *in vitro*. P-values were calculated using Kruskall-wallis tests within each feature and between different conditions.

**Figure S12. Unitig 9 shows higher nucleotide diversity.** Boxplots represent the distribution of nucleotide diversity (*π*) values calculated per 1 kbp windows in unitig 3 (Za100) and unitig 9 (Zt469). P-value was calculated using Wilcoxon rank sum test.

### Supplementary Tables

**Table S1. Summary metrics of Zt469 and Za100 genome assemblies, gene and TE annotations.**

Assembly metrices were accessed with the software Quast (Gurevich A, Saveliev V, Vyahhi N, Tesler G. 2013, Bioinformatics 29:1072–1075, https://doi.org/10.1093/bioinformatics/btt086)

**Table S2. Conditions and thresholds used for gene categories classification.**

In order to classify gene models into categories involved in host-pathogen interactions as e.g. candidates effectors and CAZymes, we used individual tools thresholds recommended by the Predector software (Jones DAB, Rozano L, Debler JW, Mancera RL, Moolhuijzen PM, Hane JK. 2021, Sci Rep 11:1–13, https://doi.org/10.1038/s41598-021-99363-0). “Small Secreted Proteins (SSPs)” category also followed the criteria described previously (Grandaubert, J., Bhattacharyya, A., & Stukenbrock, E. H. 2015, G3 Genes, Genomes, Genetics 5(7), 1323-1333, https://doi.org/10.1534/g3.115.017731) and Antismash v.6.0 (fungal version; Blin K, Shaw S, Kloosterman AM, Charlop-Powers Z, Van Wezel GP, Medema MH, Weber T. 2021, Nucleic Acids Res 49:W29–W35, https://doi.org/10.1093/nar/gkab335) was used to detected biosynthetic gene clusters (BGCs). Filtering based on these conditions and thresholds was performed independently for each gene category.

**Table S3. Summary of gene models, functional annotations and gene expression *in vitro* and *in planta* during four infection stages in Zt469.**

Functional annotations were based on several prediction tools as Predector (Jones DAB, Rozano L, Debler JW, Mancera RL, Moolhuijzen PM, Hane JK. 2021, Sci Rep 11:1–13, https://doi.org/10.1038/s41598-021-99363-0), eggnog-mapper (Huerta-Cepas J, Forslund K, Coelho LP, Szklarczyk D, Jensen LJ, Von Mering C, Bork P. 2017, Mol Biol Evol 34:2115– 2122, https://doi.org/10.1093/molbev/msx148) and Antismash v.6.0 (fungal version; Blin K, Shaw S, Kloosterman AM, Charlop-Powers Z, Van Wezel GP, Medema MH, Weber T. 2021, Nucleic Acids Res 49:W29–W35, https://doi.org/10.1093/nar/gkab335). Gene expressions *in vitro* and *in planta* are reported as the mean expression between triplicates in log2(TPM+1).

**Table S4. Summary of gene models, functional annotations and gene expression *in vitro* in Za100.**

Functional annotations were based on several prediction tools as Predector (Jones DAB, Rozano L, Debler JW, Mancera RL, Moolhuijzen PM, Hane JK. 2021, Sci Rep 11:1–13, https://doi.org/10.1038/s41598-021-99363-0), eggnog-mapper (Huerta-Cepas J, Forslund K, Coelho LP, Szklarczyk D, Jensen LJ, Von Mering C, Bork P. 2017, Mol Biol Evol 34:2115– 2122, https://doi.org/10.1093/molbev/msx148) and Antismash v.6.0 (fungal version; Blin K, Shaw S, Kloosterman AM, Charlop-Powers Z, Van Wezel GP, Medema MH, Weber T. 2021, Nucleic Acids Res 49:W29–W35, https://doi.org/10.1093/nar/gkab335). Gene expression *in vitro* is reported as the mean expression between triplicates in log2(TPM+1).

**Table S5. Summary of gene models per category in Zt469 and Za100.**

We classified the gene models into categories based on the conditions and thresholds described in Supplementary Table S2.

**Table S6. List of primers and expected amplicon sizes used in this study.**

**Table S7. Summary of genome-wide RIP signatures detected in Za100 and Zt469 isolates.** RIP-like signatures were detected using the RIPper software (Van Wyk S, Harrison CH, Wingfield BD, De Vos L, Van Der Merwe NA, Steenkamp ET. 2019, PeerJ 2019:1–18, https://doi.org/10.7717/peerj.7447). Large RIP-affected genomic regions (LRARs) were defined as genomic regions (windows) consecutively affected by RIP that are more than 4000bp in length.

**Table S8. Summary metrics of RNA-seq data in number of reads and percentage of *Zymoseptoria* genome coverage *in vitro* and *in planta*.**

**Table S9. Average expression of genes and Transposable Elements (TEs) per unitig in Zt469 during *in vitro* growth and during *in planta* infection stages.**

Gene and TE expression *in vitro* and *in planta* were calculated as the mean expression between triplicates in TPM (Transcripts Per Million). Averages per unitig and per condition were calculated as the mean expression of genes and TEs in TPM+1 divided by the total number of these features in each unitig, and finally reported as log2(TPM+1). Except for unitig 9, only unitigs syntenic to the *Z. tritici* IPO323 reference genome are shown (unitigs 0 to 20).

**Table S10. Average expression of genes and Transposable Elements (TEs) and fold change per genome compartment in Zt469 during *in vitro* growth and during *in planta* infection stages.**

Gene and TE expression *in vitro* and *in planta* were calculated as the mean expression between triplicates in TPM (Transcripts Per Million). Averages per compartment and per condition were calculated as the mean expression of genes and TEs in TPM+1 divided by the total number of these features in each compartment and finally reported as log2(TPM+1). Except for unitig 9, only unitigs syntenic to the *Z. tritici* IPO323 reference genome are considered (unitigs 0 to 20). Core compartment is composed from unitig 0 to unitig 17 and accessory compartment refers to unitigs 9, 18, 19 and 20.

**Table S11. Summary of D statistics results.**

We performed ABBA-BABA tests among four taxa (P1, P2, P3, O) in eight tree topologies to determine whether interspecific gene flow takes place. In these tests, we included the *Aegilops*-infecting *Z. tritici* isolates Zt469 (with unitig 9) or Zt436 (without unitig 9) as P1; the wheat-infecting *Z. tritici* isolates Zt565 or Zt668 as P2; and the *Z. ardabiliae* isolates Za100 (with unitig 3) or Za98 (without unitig 3) as P3. We performed all tests including the genome of the *Z. passerini* isolate Zpa796 as outgroup (O) and using a block jacknife of 15 kb.

**Table S12. List of datasets and sequencing data analyzed in this study.**

Summary of datasets generated and analyzed during this study. Metadata, accession numbers, repositories and references are provided for each *Zymoseptoria* isolate.

### Supplementary Notes

**Text S1. Additional material and methods.**

