## Supplementary material for "Interspecies hybridization as a route of accessory chromosome origin in fungal species infecting wild grasses": Fagundes_2024_Zymo_NewChr_Supplementary_Material: Supplementary_Text_S1.docx

### Supplementary Text S1 for:

**Interspecies hybridization as a route of accessory chromosome origin in fungal species infecting wild grasses**

Wagner C. Fagundes^1^*#, Mareike Möller^1,2^, Alice Feurtey^1,3,4^, Rune Hansen^1^, Janine Haueisen^1^, Fatemeh Salimi^5^, Alireza Alizadeh^6^, Eva H. Stukenbrock^1^#

^1^Environmental Genomics Group, Max Planck Institute for Evolutionary Biology, Plön & Christian-Albrechts University Kiel, Kiel, Germany

^2^Present Address: Research School of Biology, The Australian National University, Canberra, Australia

^3^ Present Address: Laboratory of Evolutionary Genetics, Institute of Biology, University of Neuchâtel, Neuchâtel, Switzerland

^4^ Present Address: Plant Pathology Group, Institute of Integrative Biology, ETH Zürich, Zürich, Switzerland

^5^Department of Plant Protection, College of Agriculture and Natural Resources, University of Tehran, Karaj, Iran

^6^Department of Plant Protection, Faculty of Agriculture, Azarbaijan Shahid Madani University, Tabriz, Iran

#Address correspondence to Eva H. Stukenbrock, and Wagner C. Fagundes,

*Present address: Adaptive Evolution of Filamentous Plant Pathogens, Max Planck Institute for Biology, Tübingen, Germany

**Material and methods**

**ChIP- and RNA-sequencing of *in vitro* cultures**

We used three biological replicates of the *Z. tritici* isolate Zt469 and *Z. ardabiliae* isolate Za100 grown *in vitro* for chromatin immunoprecipitation followed by sequencing (ChIP-seq) and RNA-sequencing. Cells from the same biological replicate were used for both RNA and ChIP DNA extractions. Fungal spores were grown on YMS plates for three to four days, harvested and resuspended in 6 mL of 1x PBS (137 mM NaCl, 2.7 mM KCl, 10 mM Na2HPO4, 1.8 mM KH2PO4). For RNA extraction, 1 mL of resuspended cells were centrifuged, ground in liquid nitrogen, and total RNA was extracted using TRIzol (Invitrogen, Karlsruhe, Germany) following the manufacturer’s instructions. Cleaned (DNAse-treated) RNA samples were sent to Admera Health (South Plainfield, NJ, USA) for library preparation. For ChIP, the remaining 5 mL of resuspended cells were crosslinked with 0.5% formaldehyde for 15 min at room temperature and quenched by adding 150 μL of 2.5 M glycine. Chromatin immunoprecipitation was performed as previously described (1, 2) and antibodies against H3K4me2 (#07–030, Merck Millipore), H3K9me3 (#39161, Active Motif) and H3K27me3 (#39155, Active Motif) were used. ChIP-seq libraries were prepared with a modified version of the Next Ultra II DNA Library Prep Kit for Illumina (#E7645S, New England Biolabs). Sequencing of ChIP-seq and RNA-seq samples was performed on an Illumina HiSeq 3000, obtaining paired-end reads of 150 nt by Admera Health (South Plainfield, NJ, USA).

**Pulsed-Field Gel Electrophoresis (PFGE) and Southern blot analyses**

For PFGE analyses, we used chromosomal DNA of *Hansenula wingei* (Bio-Rad, Munich, Germany) as standard size marker for mid-size chromosomes (1 to 3.1 Mb). PFGE run settings were followed as described previously (3). Gels were stained with ethidium bromide solution (1 mg/ml ethidium bromide in H2O) for 30 min and chromosomal bands were detected with the GelDocTM XR+ system (Bio-Rad). We performed Southern blot analysis as described previously (4) using a DIG (digoxigenin)-labeled probe generated by the PCR digoxigenin labeling Mix (Roche, Mannheim, Germany) following the manufacturer’s instructions. Probe primers were designed with Primer-BLAST (5) and Geneious v.2020.1.2 software (https://www.geneious.com/home/). List of primers used can be found at Supplementary Table S6.

**Introgression analyses**

In order to compute the ABBA-BABA tests, whole-genome paired-end sequencing reads (2x150bp) were first trimmed using Trimmomatic v 0.39 (6) with the following parameters: LEADING:20 TRAILING:20 SLIDINGWINDOW:5:20 MINLEN:50. Trimmed reads were then aligned to the *Z. tritici* IPO323 reference genome (7) using the short-read aligner bwa-mem v. 0.7.17 (8). For these tests, we only aligned reads to the core genome (chromosomes 1 to 13) of *Z. tritici* to avoid biases due to presence-absence variation of accessory chromosomes. Alignments were converted to binary files (.bam) and sorted using SAMtools v. 1. 13 (9). Read duplicates and read groups were determined using Picard tools v. 2.26.2 (<https://broadinstitute.github.io/picard/>) and alignment files were then finally used as input in the “D-stat” module from the ANGSD package (Analysis of Next Generation Sequencing Data; <https://www.popgen.dk/angsd/index.php/Abbababa>; (10). We used a block size of 15kb (-blockSize 15000) to account for possible linkage disequilibrium (LD) blocks based on previous reports of LD decay in *Z. tritici* populations (11–13) and only kept sites where all individuals have data (-minInd 4) with a minimum mapping quality score of 20 (-minMapQ 20).

**Variant calling**

In order to analyze nucleotide diversity per chromosome, we generated Variant Calling Format (VCF) files using short-read population data of *Aegilops*-infecting *Z. tritici* and *Z. ardabiliae* isolates obtained from previous studies (13–16). Only the 11 *Aegilops*-infecting *Z. tritici* and 3 *Z. ardabiliae* isolates with reads mapped to unitig 9 (in Zt469) and unitig 3 (in Za100), respectively, were used for these analyses. First, we trimmed whole-genome paired-end sequencing reads (2x150bp) using Trimmomatic v 0.39 (6) with the following parameters: LEADING:20 TRAILING:20 SLIDINGWINDOW:5:20 MINLEN:50. Trimmed reads were then aligned to the *Aegilops*-infecting *Z. tritici* Zt469 (for *Aegilops*-infecting *Z. tritici* isolates) and *Z. ardabiliae* Za100 (for *Z. ardabiliae* isolates) reference genomes using the short-read aligner bwa-mem v. 0.7.17 (8). Conversion, sorting, merging, and indexing of alignment files were performed using SAMtools v. 1. 13 (9). PCR duplicates and read groups were determined using Picard tools v. 2.26.2 (https://broadinstitute.github.io/picard/). Single Nucleotide Polymorphism (SNP) calling, genotyping and variant filtration were performed using the Genome Analysis Toolkit (GATK) v. 4.1.4.1 (17) for each species individually. SNPs were hard-filtered using the GATK VariantFiltration and SelectVariants tools. We first filtered low depth genotypes using the following parameters: --genotype-filter-expression "DP < 3", --set-filtered-genotype-to-no-call and --genotype-filter-name "low_depth". Then, the following filter conditions were applied: DP > 1000.0 (for *Aegilops*-infecting *Z. tritici*) and DP > 2000.0 (for *Z.ardabiliae*) ; QD < 20.0; MQ < 50.0; FS > 20.0; ReadPosRankSum, MQRankSum, and BaseQRankSum between -2 and 2. At last, we kept only biallelic SNPs with a genotyping rate of ≥ 90% for each species using VCFtools v 0.1.13 (18).
