## Supplementary figures and images for "Interspecies hybridization as a route of accessory chromosome origin in fungal species infecting wild grasses"

### FigureS1.tiff

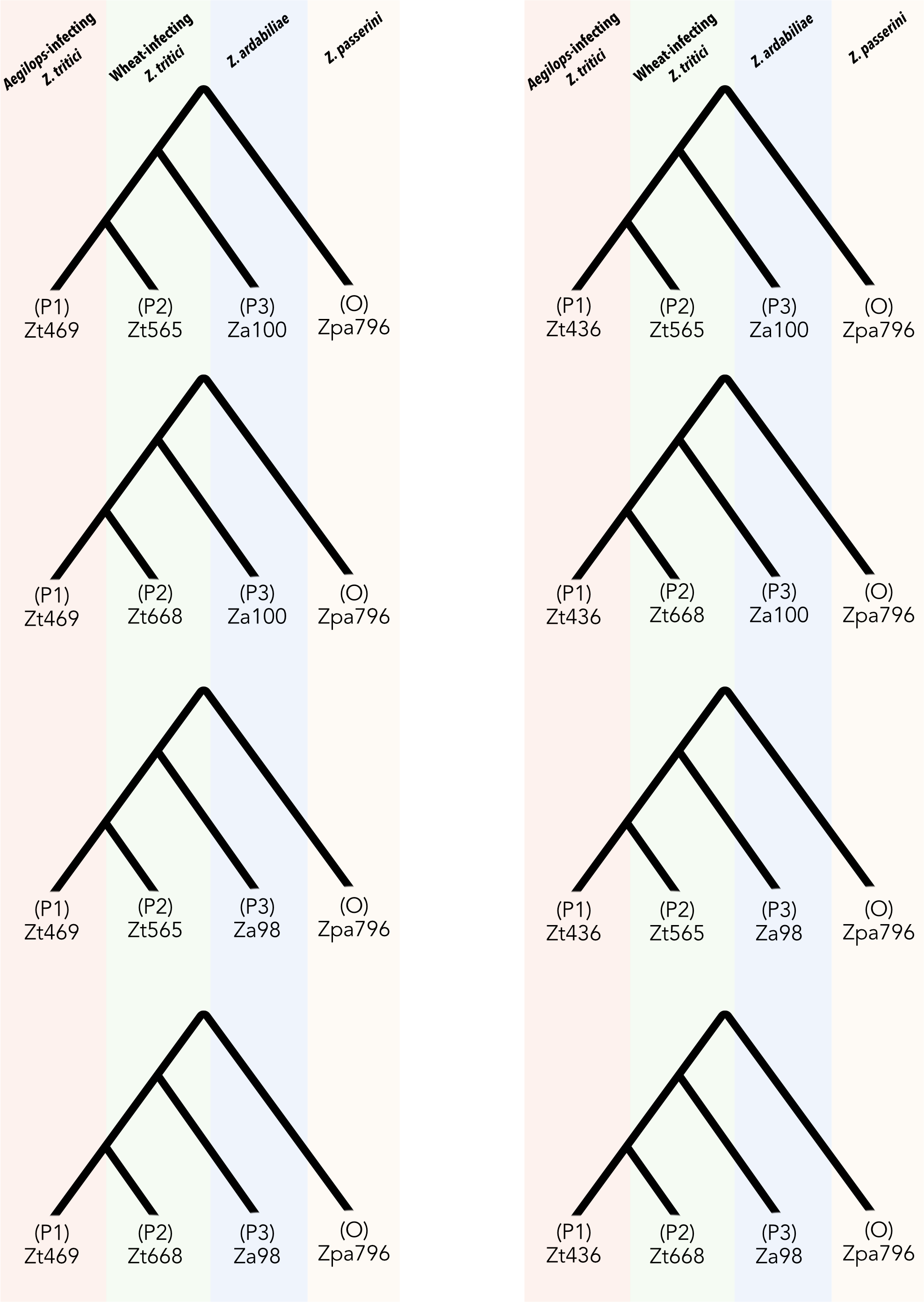

### FigureS2.tiff

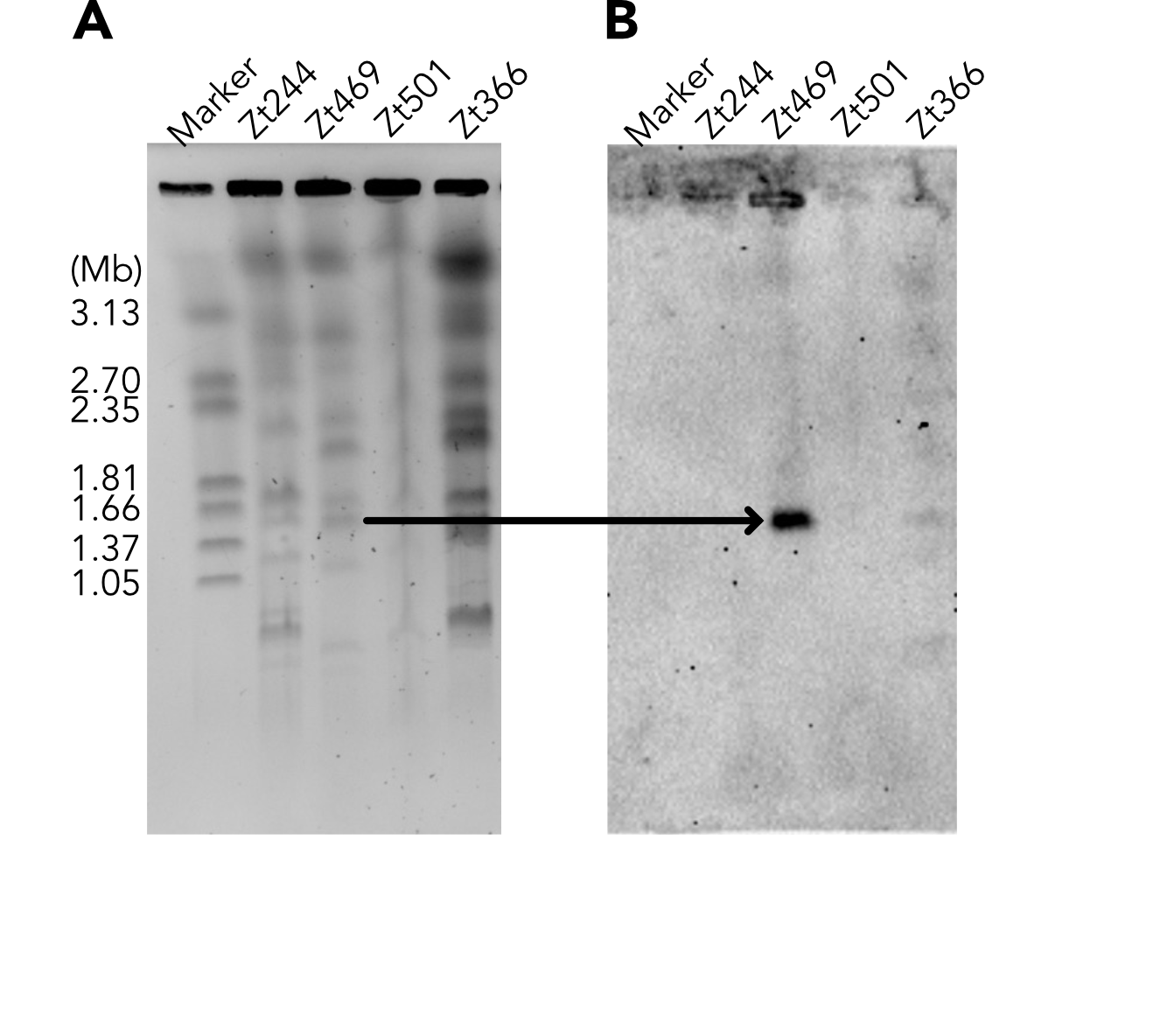

### FigureS3.tiff

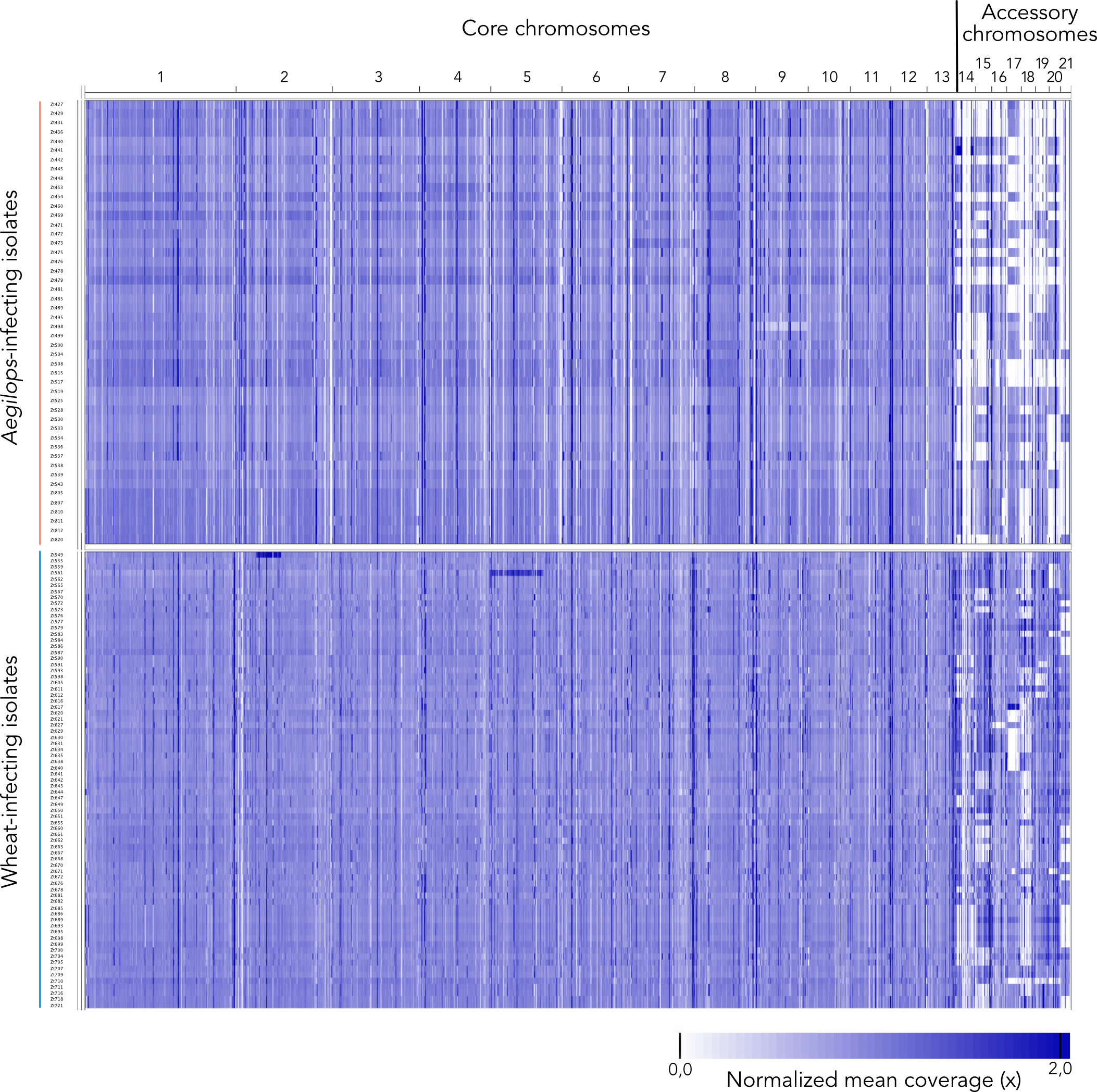

### FigureS4.tiff

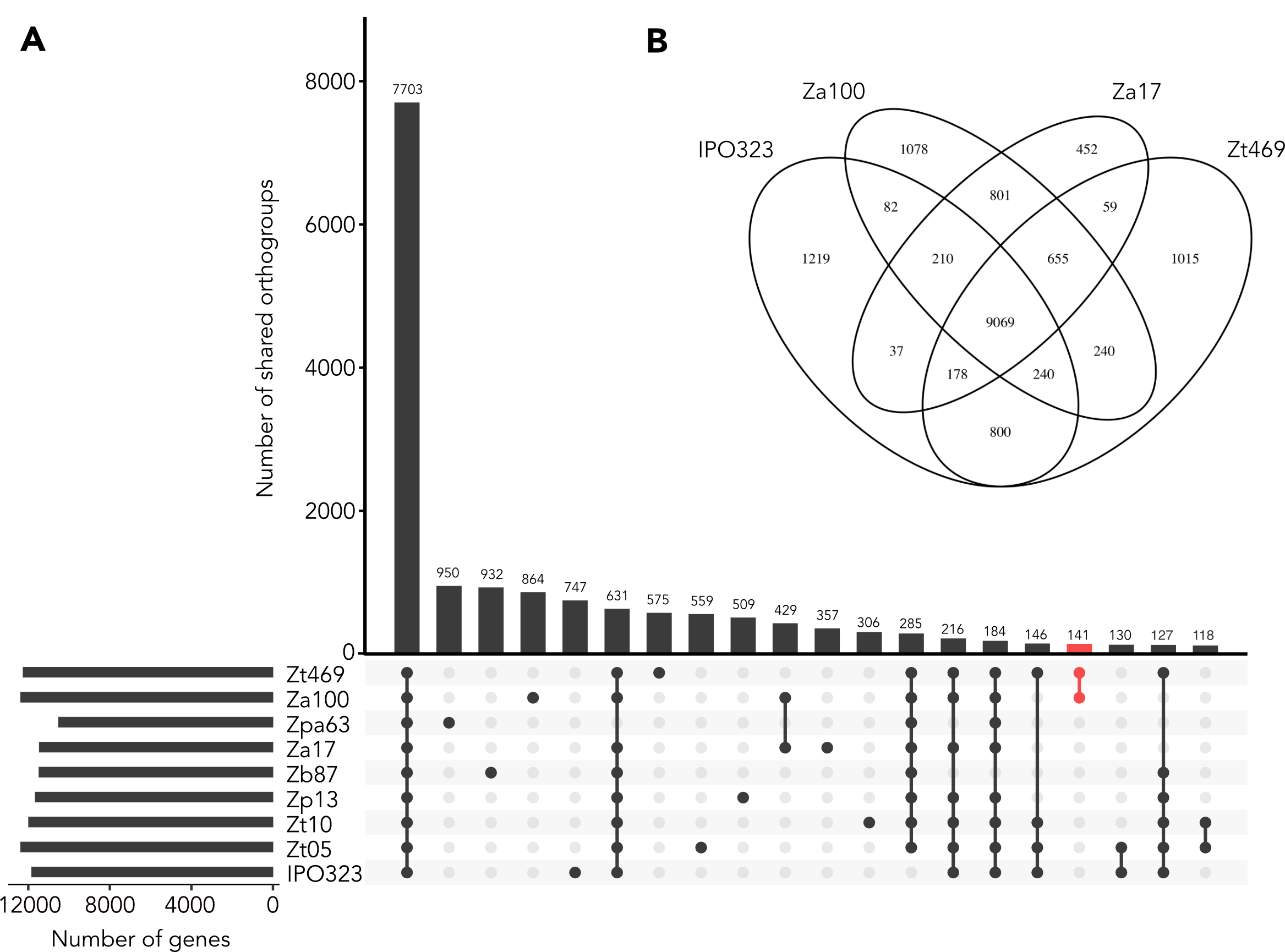

### FigureS5.tiff

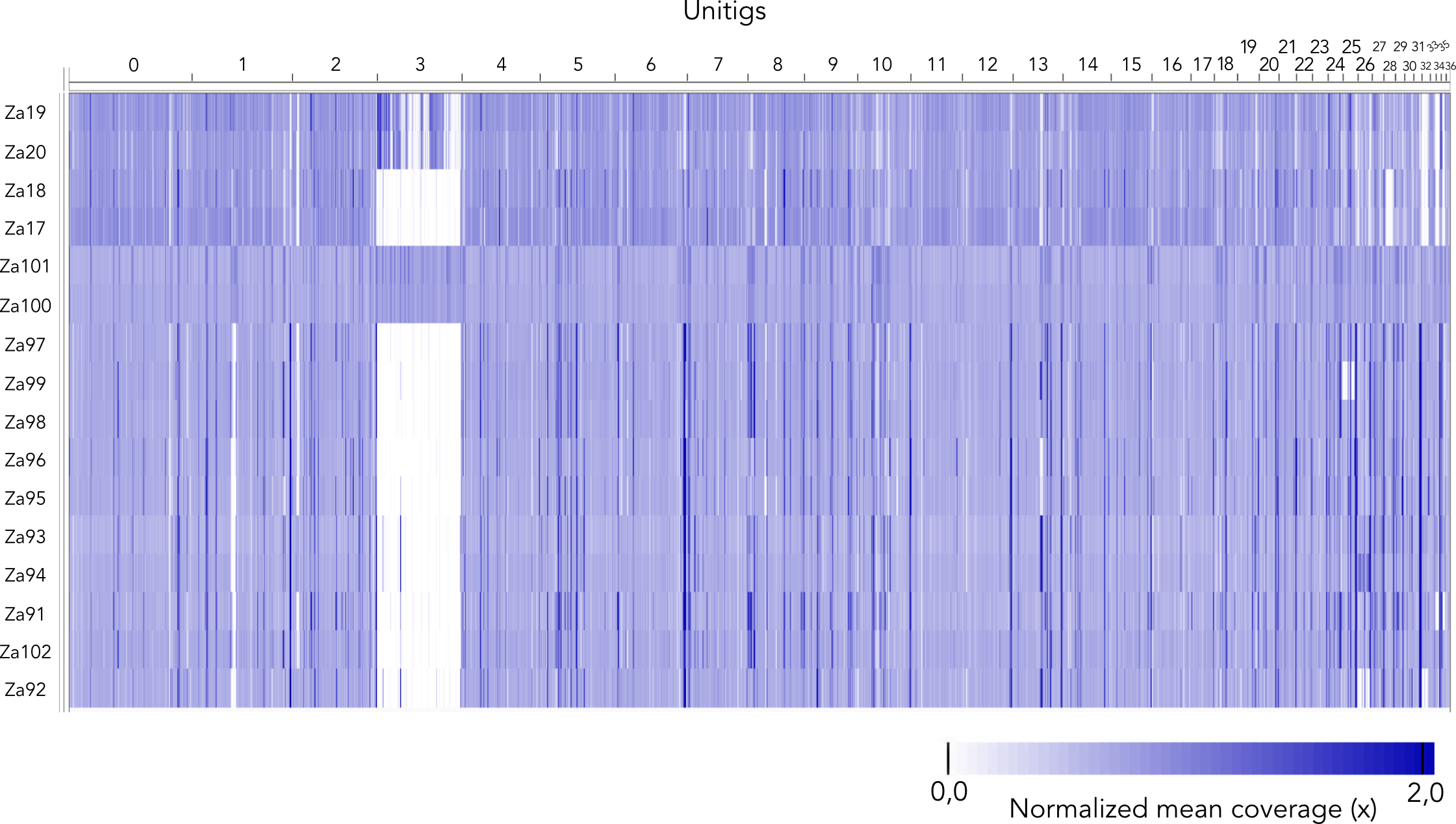

### FigureS6.tiff

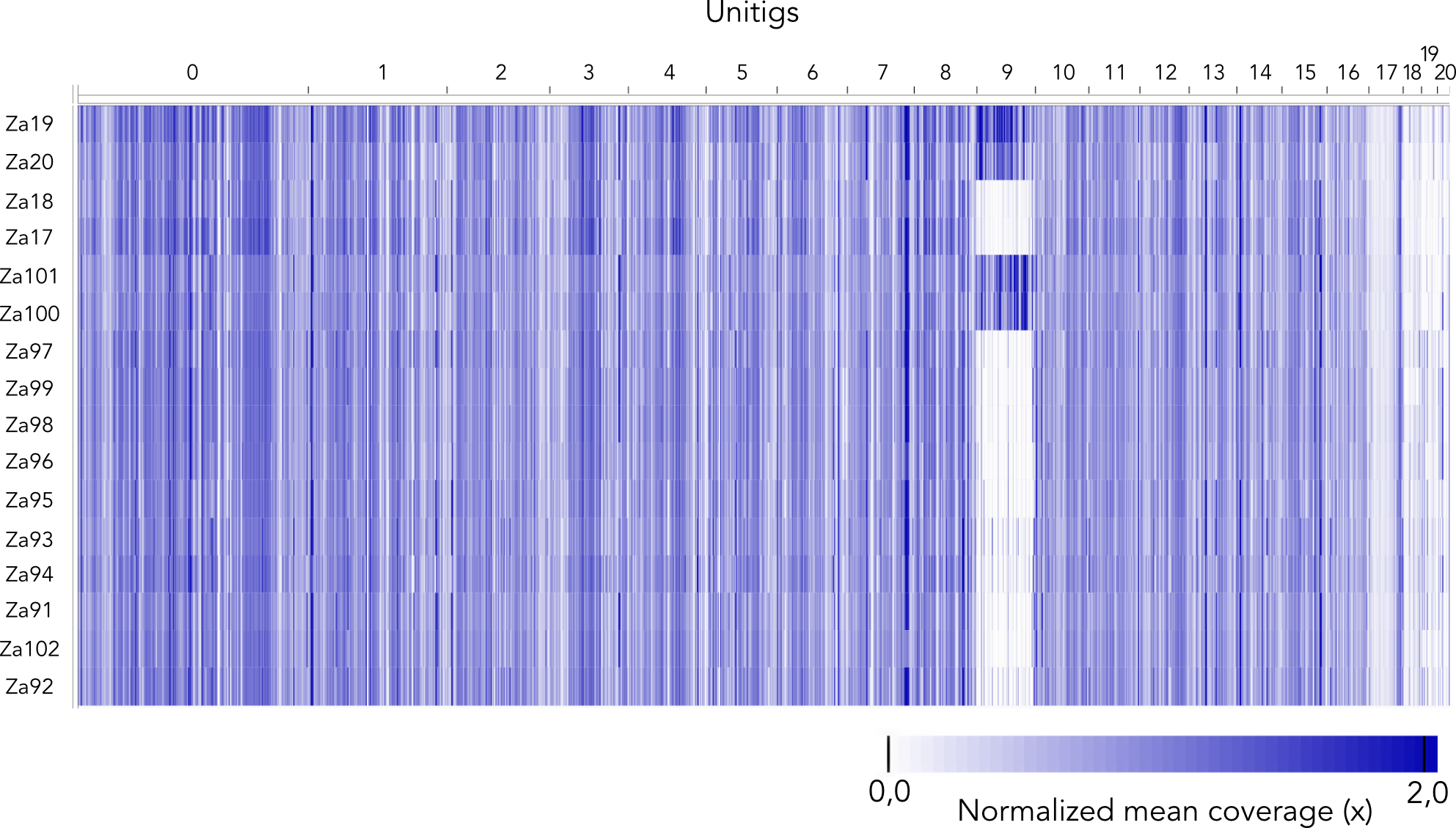

### FigureS7.pdf

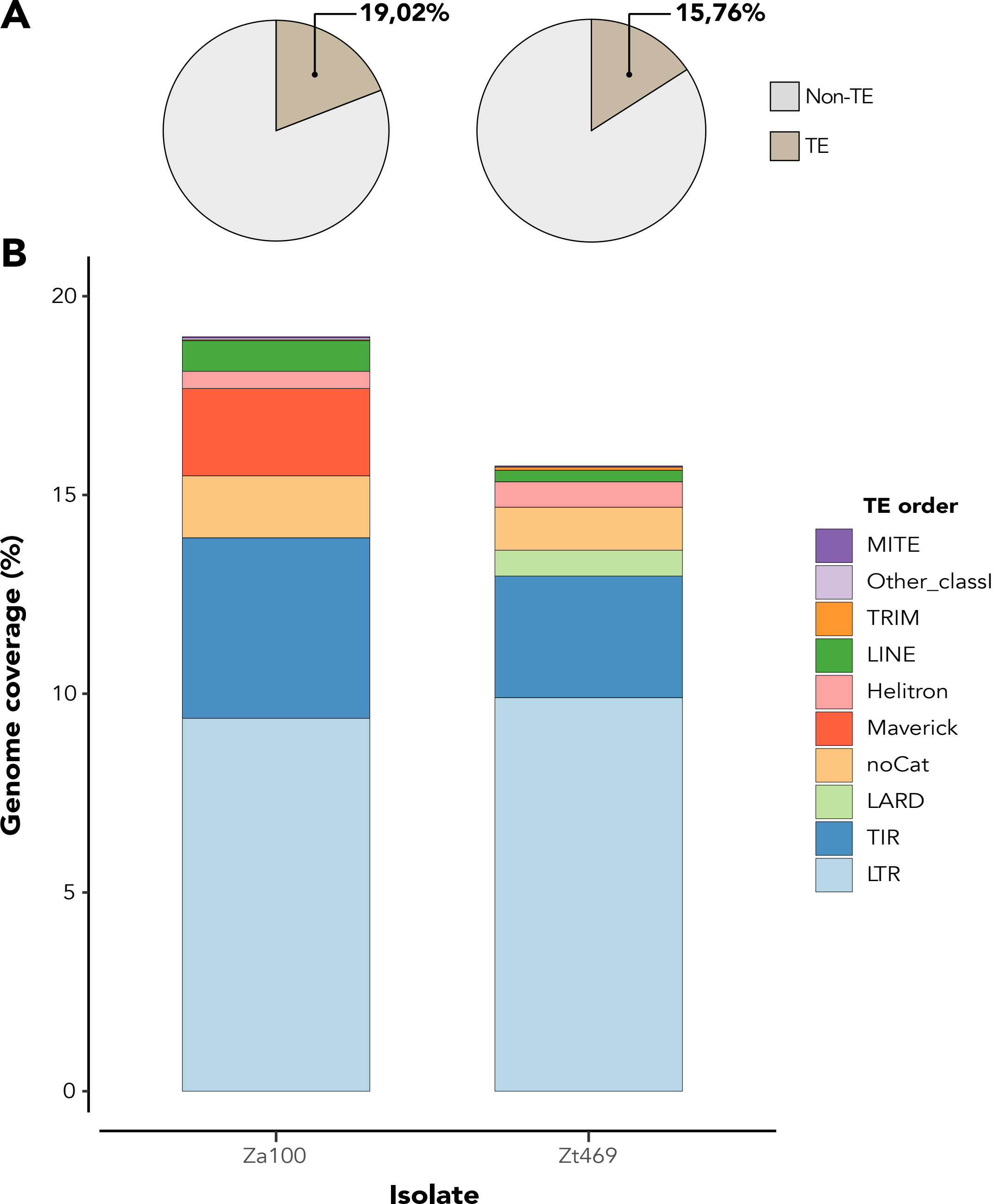

### FigureS8.pdf

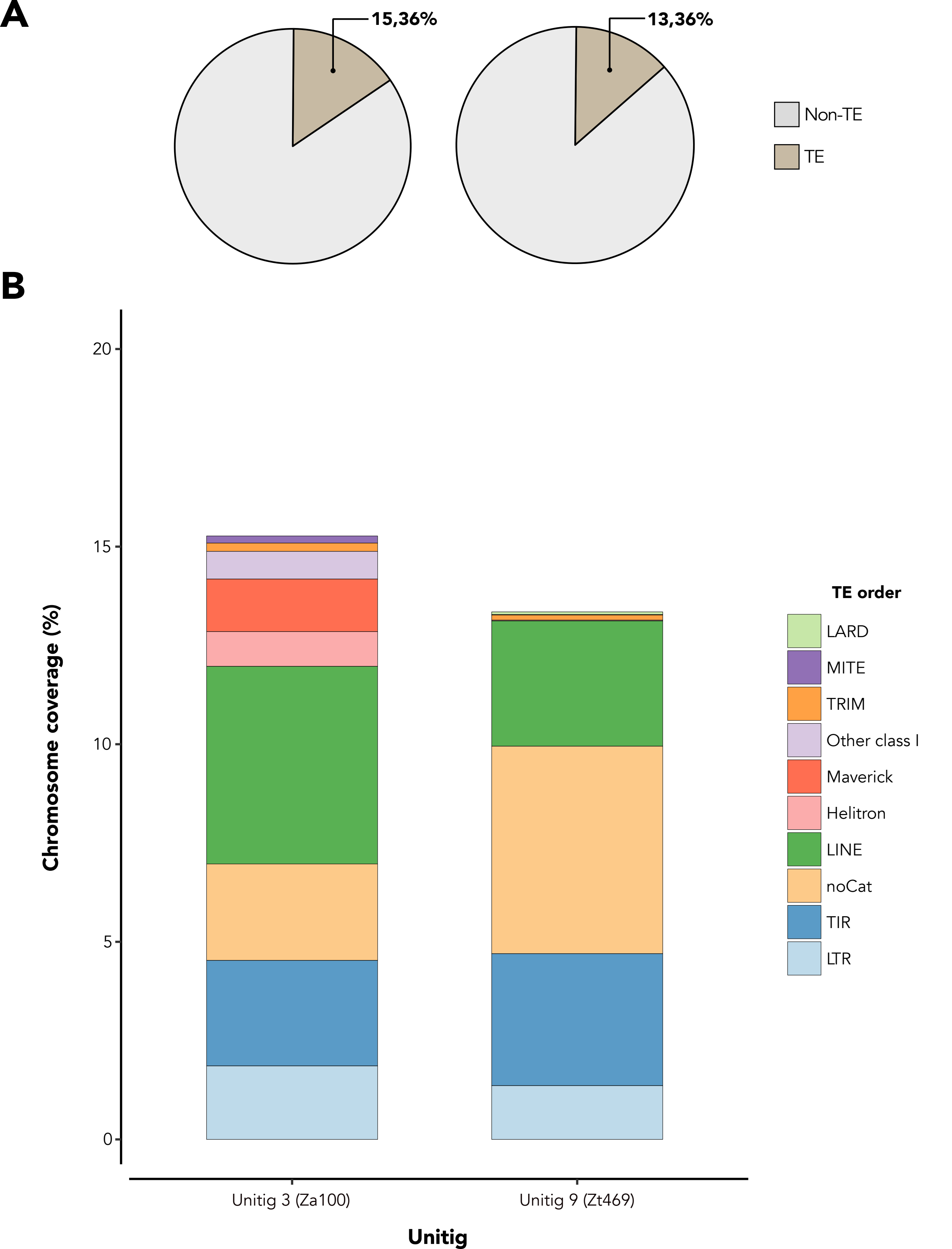

### FigureS9.tiff

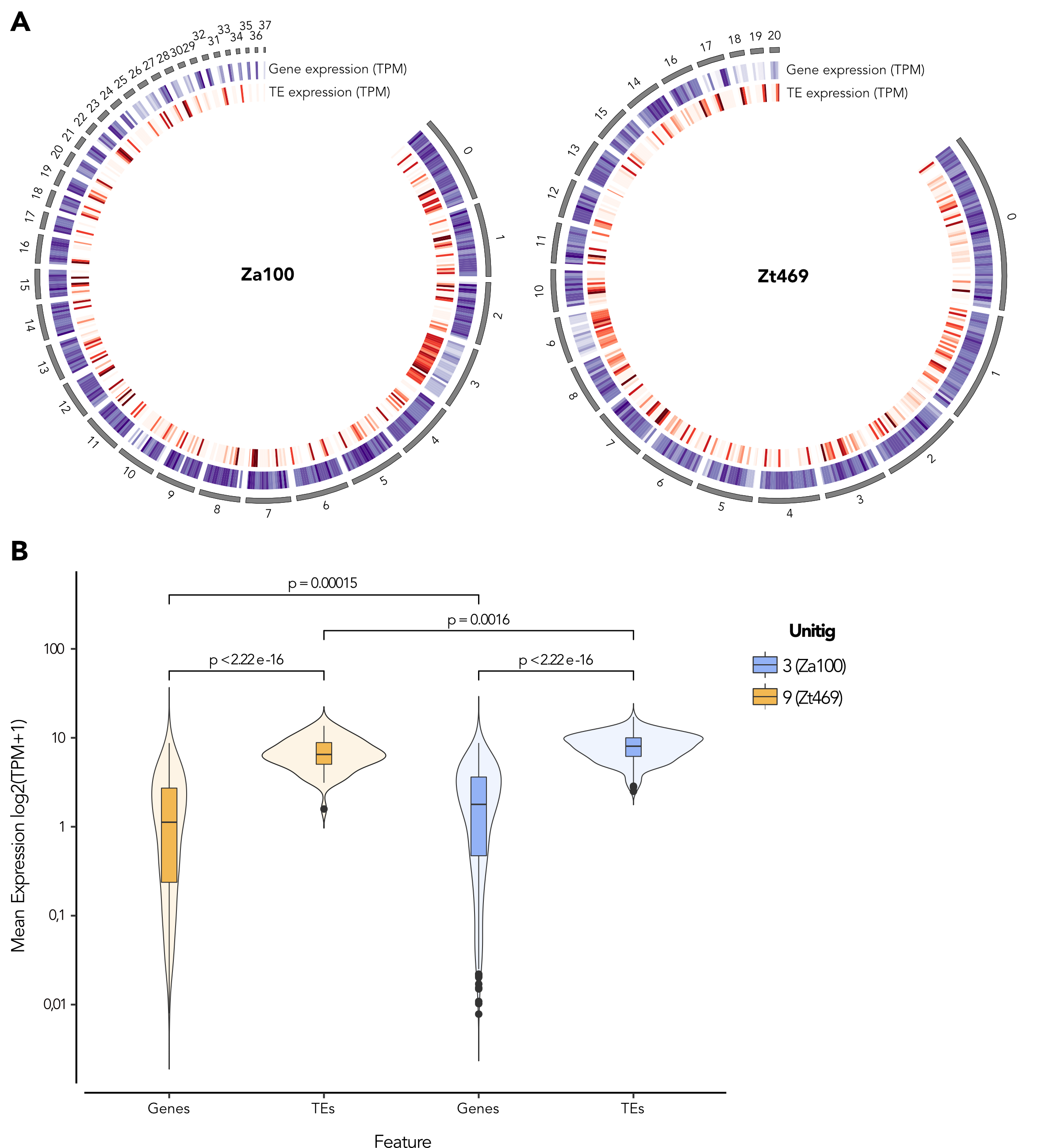

### FigureS10.tiff

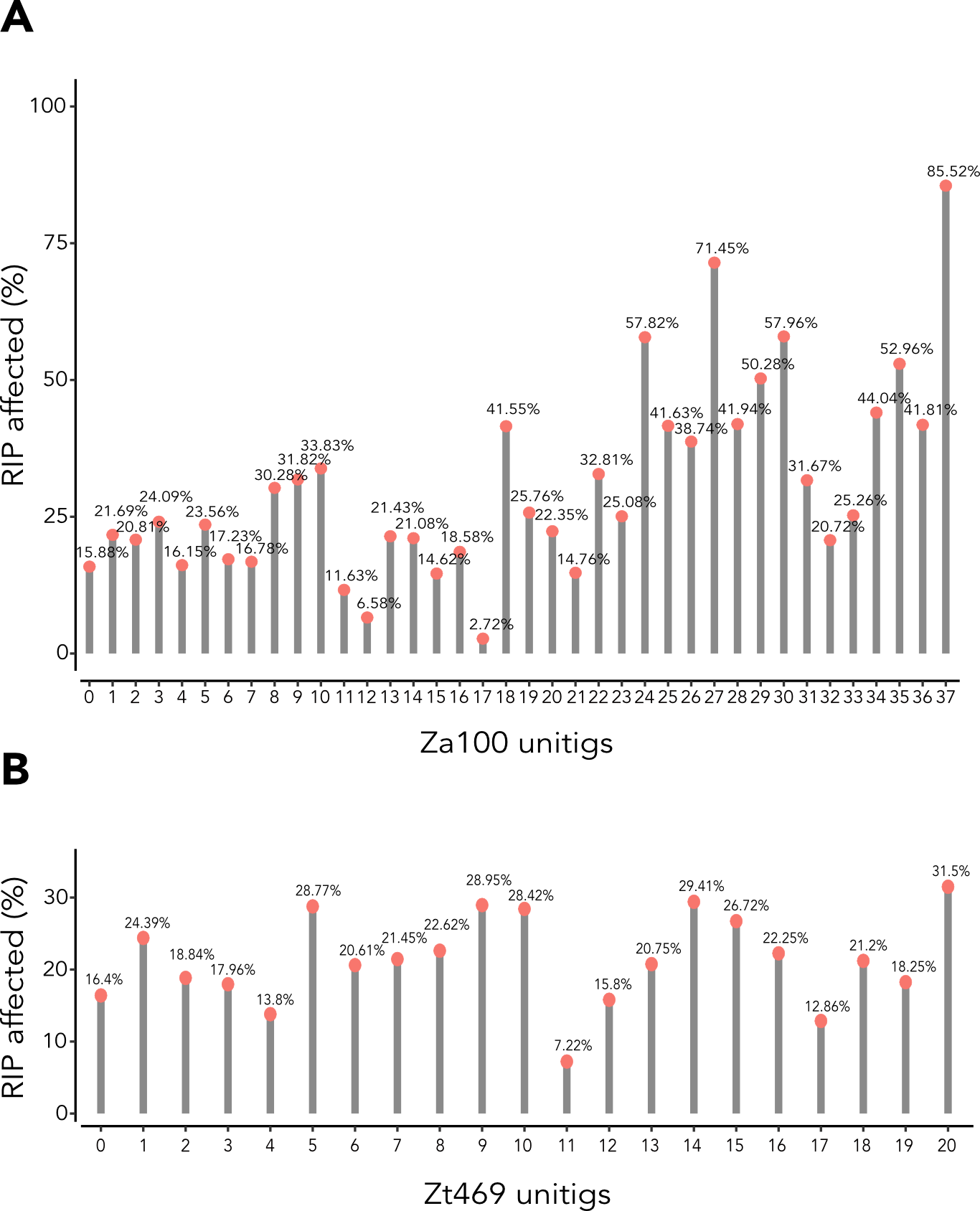

### FigureS11.tiff

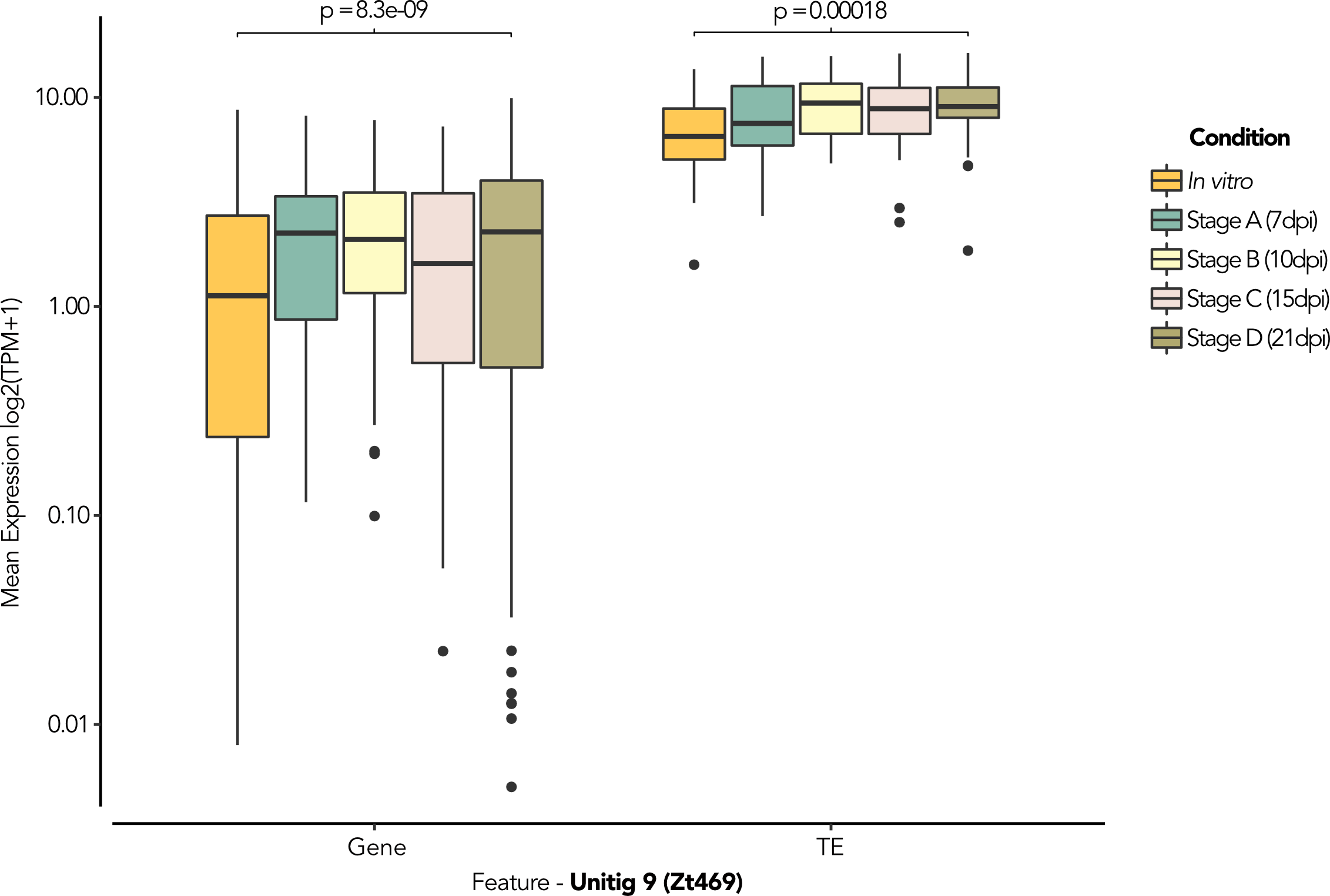

### FigureS12.tiff

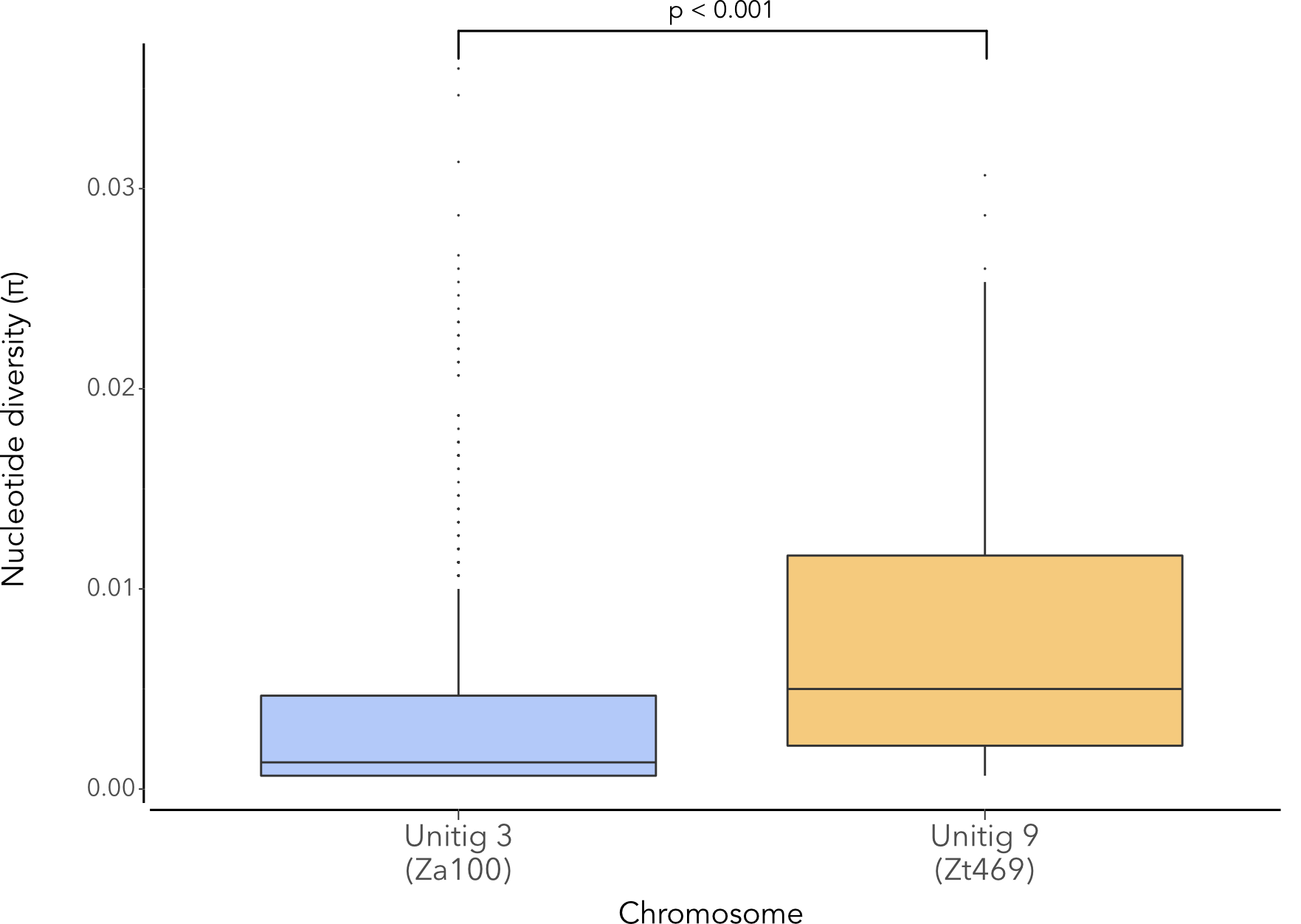
